# Discovery and Preclinical Validation of a Clinically Optimized Mitochondrial Complex I Modulator for Alzheimer’s Disease

**DOI:** 10.64898/2026.04.10.717554

**Authors:** Sergey Trushin, Thi Kim Oanh Nguyen, Mark Ostroot, Alexander Galkin, Toshihiko Nambara, Wenyan Lu, Takahisa Kanekiyo, Graham Johnson, Eugenia Trushina

## Abstract

Alzheimer’s disease (AD) is characterized by diminished capacity to mount adaptive cellular stress responses required to maintain energy homeostasis and proteostasis. An emerging therapeutic strategy is to restore adaptive stress responses by inducing mild energetic stress through inhibition of mitochondrial complex I (mtCI). However, pharmacological inhibition of the respiratory chain has remained challenging, as it can induce bioenergetic failure rather than beneficial signaling. Here, we describe C273, a brain-penetrant small molecule that delivers controlled, weak attenuation of mtCI activity to therapeutically restore endogenous adaptive stress pathways. This work establishes a first-in-class mechanism in which calibrated activation of multifaceted adaptive mechanisms enhances cellular resilience, rather than impairing mitochondrial function. Structure-activity relationship optimization yielded a compound with high potency against Aβ-induced cellular toxicity, strong selectivity for mtCI, and favorable drug-like properties. C273 demonstrated excellent oral bioavailability, metabolic stability in mouse, rat, and human microsomes, minimal CYP liabilities, and a clean ancillary pharmacology profile in the Eurofins CEREP44 panel. *In vivo*, C273 readily crosses the blood-brain barrier and activates AMP-activated protein kinase (AMPK), initiating a coordinated hormetic response characterized by enhanced antioxidant defenses, suppression of inflammatory signaling, induction of autophagy, and increased mitochondrial biogenesis and turnover. Genetic deletion of AMPKα1/α2 abolished these responses, establishing AMPK as a critical mediator of C273 activity. Pharmacological competition experiments further confirmed the target, as pretreatment with non-toxic concentrations of rotenone blocked C273 interaction with the quinone-binding site of mtCI and eliminated its neuroprotective effects. Repeated oral administration of C273 (20-80 mg/kg/day) to wild-type mice for one month produced no detectable cardiac or hepatic toxicity, indicating a favorable *in vivo* safety margin. Importantly, C273 activated these mechanisms and reduced Aβ and p-Tau levels in induced pluripotent stem cell-derived cerebral organoids from patients with sporadic AD. Collectively, these results establish controlled mtCI modulation as a therapeutic strategy and position C273 as a promising disease-modifying candidate for AD.

## Introduction

Alzheimer’s disease (AD) is a progressive neurodegenerative disorder characterized by amyloid-β (Aβ) accumulation, tau hyperphosphorylation, synaptic dysfunction, and neuroinflammation^1^. Although therapeutic strategies have historically focused on reducing amyloid burden, recent evidence indicates that AD is driven by multiple interconnected mechanisms, including inflammation, impaired autophagy, oxidative stress, and mitochondrial dysfunction, thereby necessitating polypharmacological approaches^2–6^. Impaired oxidative phosphorylation (OXPHOS) machinery, elevated levels of reactive oxygen species (ROS), reduced mitochondrial biogenesis and defective mitophagy are consistently observed in AD patient brains and in transgenic mouse models of AD^7^. Mitochondrial signaling is tightly integrated with oxidative stress, inflammation, and epigenetic regulation, positioning mitochondria as central hubs that coordinate multiple disease-relevant pathways. Notably, mitochondrial dysfunction emerges early, often preceding overt neurodegeneration, and can potentiate both Aβ and tau pathology. Together, these features establish mitochondrial pathways as mechanistically upstream, highly integrative, and still underexploited therapeutic targets^8–10^.

Mitochondrial complex I (mtCI; NADH:ubiquinone oxidoreductase) is the only entry point for NADH-derived electrons into the electron transport chain (ETC)^11^. It is a key component for proton motive force generation and the main regulator of cellular redox and NAD⁺/NADH balance, as well as ROS-mediated signaling^12,13^. Through these functions, mtCI integrates mitochondrial bioenergetic status with broader cellular stress responses and disease processes^14–16^. While complete inhibition of mtCI by toxins such as rotenone or piericidin A disrupts mitochondrial respiration and induces neurotoxicity^17^, emerging evidence suggests that weak, reversible modulation of mtCI activity can instead activate adaptive cellular stress responses^18^. In particular, modest energetic shifts produced by mild mtCI inhibition, similar to those induced by physiological stresses such as exercise or caloric restriction, or by metformin, activate AMP-activated protein kinase (AMPK)^19,20^. AMPK functions as a central metabolic regulator controlling mitochondrial biogenesis, autophagy, and inflammatory signaling^20,21^. Activation of this pathway promotes PGC-1α-dependent mitochondrial remodeling^22^, enhances antioxidant defenses^23^, and suppresses NF-κB-mediated inflammatory signaling pathways^24^, which are dysregulated in AD.

Consistent with this concept, we previously demonstrated that mtCI represents a druggable target whose controlled modulation can activate neuroprotective signaling pathways and improve pathology in multiple AD mouse models exhibiting Aβ (APP/PS1, APP, PS1, 5xTgAD) and tau (3xTgAD) pathologies^25–33^. The tricyclic pyrone compound CP2 provided proof of concept that mild mtCI modulation can restore mitochondrial homeostasis, ameliorate cognitive and molecular deficits, and block the ongoing neurodegeneration in AD models^10,34^. Subsequent structure-activity relationship (SAR) optimization yielded a new series of molecules where a new lead compound C458 had improved target selectivity and *in vivo* efficacy^27^. However, additional medicinal chemistry optimization was required to obtain a compound with pharmacokinetic, safety, and drug-like properties suitable for clinical development.

Here, we report the rational design and preclinical characterization of C273, a clinically optimized weak mtCI inhibitor derived from CP2 and C458 through iterative SAR studies. Compounds were advanced through an integrated drug discovery workflow designed to improve potency, selectivity, safety, and translational potential^27^. Screening included evaluation of cellular protection against Aβ-induced toxicity, assessment of cytochrome P450 and hERG liabilities, metabolic stability in mouse and human microsomes, pharmacokinetic profiling, and measurement of blood-brain barrier (BBB) penetration. This strategy led to the identification of C273, a BBB-penetrant weak mtCI modulator that activates adaptive stress signaling. C273 confers neuroprotection in cellular models of AD and reduces Aβ and p-Tau accumulation in cerebral organoids derived from patients with sporadic AD.

## Results

### Design and synthesis of C273

C273 was developed through rational optimization of the previously reported lead compound C458^27^ and the tool compound CP2^26,28,33–35^. The goal of this design iteration was to generate a clinically tractable candidate with improved absorption, distribution, metabolism, and excretion (ADME) and pharmacokinetic (PK) properties while maintaining mtCI target engagement, mechanism of action and efficacy in preclinical models (**Fig. 1a**).

**Fig. 1:**
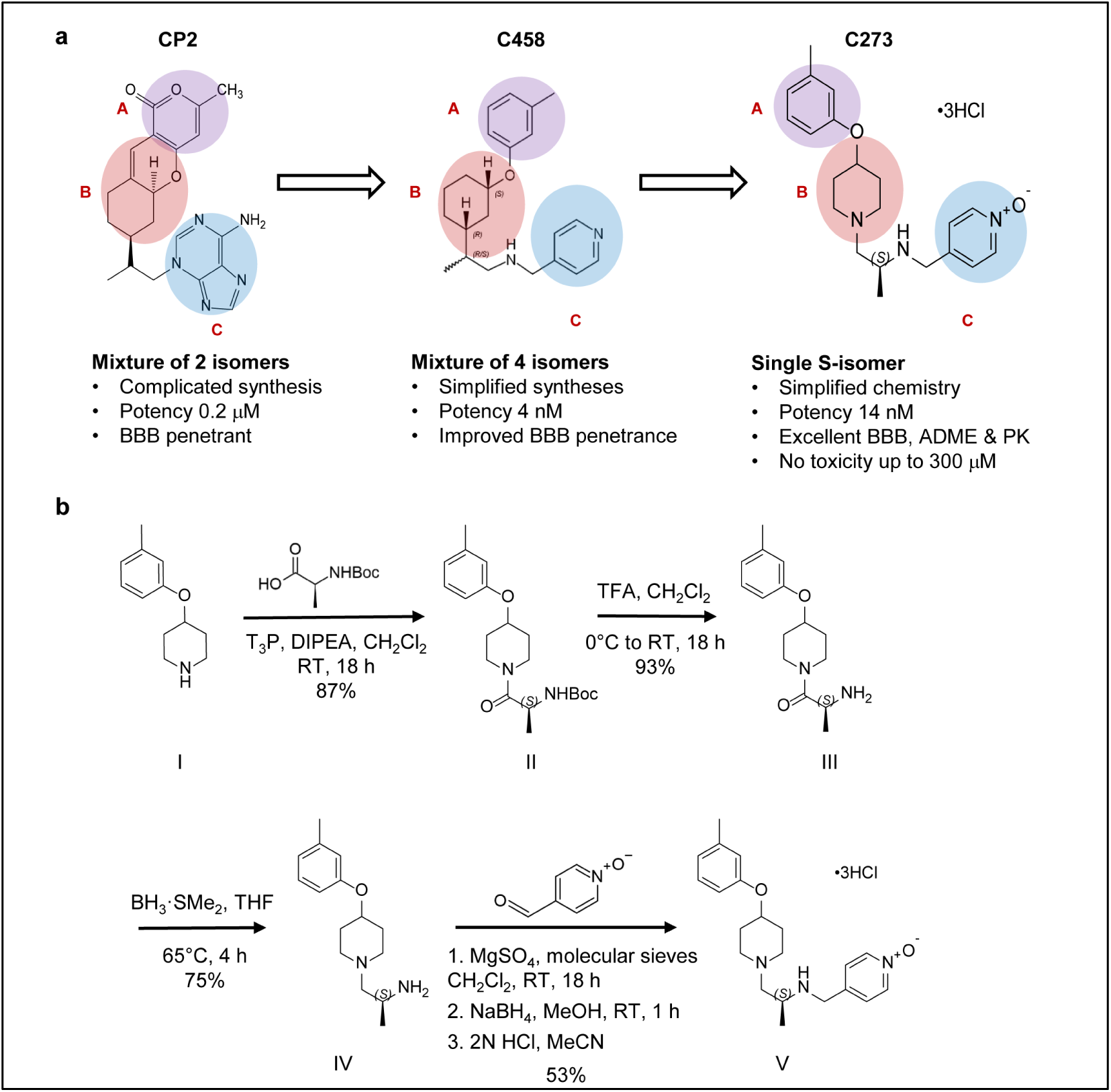
Rational design and synthesis of the mtCI modulator C273. **a** Structure-guided optimization of the lead compounds C458 and CP2. Targeted modifications in A, B, and C regions were introduced to improve pharmacological and ADME properties while preserving mtCI target engagement. **b** Synthetic route to C273. The design strategy reduced stereochemical complexity and enhanced drug-like properties, yielding the single S-enantiomer C273 (compound V) with an overall yield of 53%.

Medicinal chemistry efforts focused on reducing stereochemical complexity while preserving the structural determinants required for mtCI modulation. The resulting compound, C273 ((S)-4-(((1-(4-(m-tolyloxy)piperidin-1-yl)propan-2-yl)amino)methyl)pyridine 1-oxide), was synthesized as a single S-enantiomer (**Fig. 1b**, compound V) with an overall yield of 53%. Relative to C458, C273 incorporates a modified piperidine scaffold that reduces molecular chirality and enables a more efficient and stereoselective synthetic route. In addition, the terminal aromatic substituent (ring C) was replaced with a pyridine N-oxide moiety, a modification designed to mitigate cytochrome P450 (CYP) and voltage-gated potassium channel hERG liabilities while improving microsomal stability, plasma protein binding, and aqueous solubility.

Consistent with these design objectives, C273 exhibited favorable physicochemical properties compatible with CNS drug development (**Table 1**)^36^. Compared with CP2 and C458, C273 maintained balanced lipophilicity (calculated logP = 1.1), substantially lower than that of C458 (logP = 4.62), reducing the likelihood of nonspecific binding and off-target interactions. C273 has a total polar surface area of 51.4 Å^2^, well below the threshold generally associated with poor BBB permeability, while remaining higher than that of C458, consistent with improved solubility without compromising permeability. C273 also satisfies Lipinski’s rule of five with no violations and meets Veber criteria for oral bioavailability, while maintaining a molecular weight within the preferred range for CNS-active compounds (355.48 Da). The balanced hydrogen-bond donor/acceptor profile and moderate number of rotatable bonds further support its favorable ADME characteristics. Together, these features position C273 within a CNS-relevant drug-like chemical space while preserving the structural attributes required for selective, weak modulation of mtCI activity. The SAR studies indicated that modification of the terminal aromatic substituent enabled tuning of lipophilicity and metabolic stability while preserving the core features required for mtCI target engagement.

**Table 1.** Drug-like properties of CP2, C458 and C273.

|  | Optimal | CP2 | C458 | C273 |
| --- | --- | --- | --- | --- |
| Calculated log P | <5 | 1.86 | 4.62 | 1.1 |
| Total polar surface area | <100 Å<sup>2</sup> | 105.15 Å<sup>2</sup> | 34.15 Å<sup>2</sup> | 51.4 Å<sup>2</sup> |
| H-bond acceptors | <10 | 8 | 3 | 5 |
| H-bond donors | <5 | 2 | 1 | 1 |
| Rotatable bonds | <10 | 3 | 7 | 7 |
| Lipinski's rule of 5 violations | <1 | <1 | <1 | <1 |
| Passed both Veber criteria | yes | yes | yes | yes |
| Molecular weight | <450 | 393.4 | 338.5 | 355.48 |

### C273 retains protection against Aβ-induced toxicity with minimal intrinsic cytotoxicity

To evaluate the safety and efficacy of C273 as a weak mtCI modulator, we used a previously established drug-discovery screening funnel^27^ that assesses cytotoxicity and protection against Aβ-induced cellular toxicity across a broad concentration range (**Fig. 2a).** Compound activity was evaluated in the human neuroblastoma MC65 cells, a well-established model of intracellular Aβ toxicity. MC65 cells conditionally express the C99 fragment of amyloid precursor protein (APP) under the control of a tetracycline (Tet)-regulated promoter^37^. Upon tetracycline withdrawal (Tet-Off), C99 is expressed and processed by γ-secretase to generate Aβ peptides, leading to cell death within approximately three days (**Fig. 2b**, #). This is associated with oxytosis, ferroptosis, and mitochondrial dysfunction, processes relevant to AD pathogenesis, and provides excellent phenotypic assays for drug discovery^38^. In contrast, in the presence of tetracycline (Tet-On), C99 expression is suppressed, allowing assessment of compound cytotoxicity independently of Aβ production. C273 protected MC65 Tet-Off cells from Aβ-induced toxicity with an EC_50_ of 14.2 ± 1.8 nM (**Fig. 2b**, red curve). In the same assay, the efficacy of CP2 and C458 against Aβ toxicity was 200 nM and 4 nM, respectively^27,35^. C273 showed no detectable cytotoxicity across a broad concentration range (2 nM - 100 μM) in MC65 Tet-On cells **(Fig. 2b**, black curve) and in HepG2 human hepatic cells at concentrations up to 100 µM (**Fig. 2c**). These findings indicate that C273 retains robust neuroprotective activity in the MC65 phenotypic assay while exhibiting minimal intrinsic cytotoxicity. Notably, C273 preserved potency at nanomolar concentration while incorporating structural changes designed to improve developability and CNS drug-like properties.

**Fig. 2:**
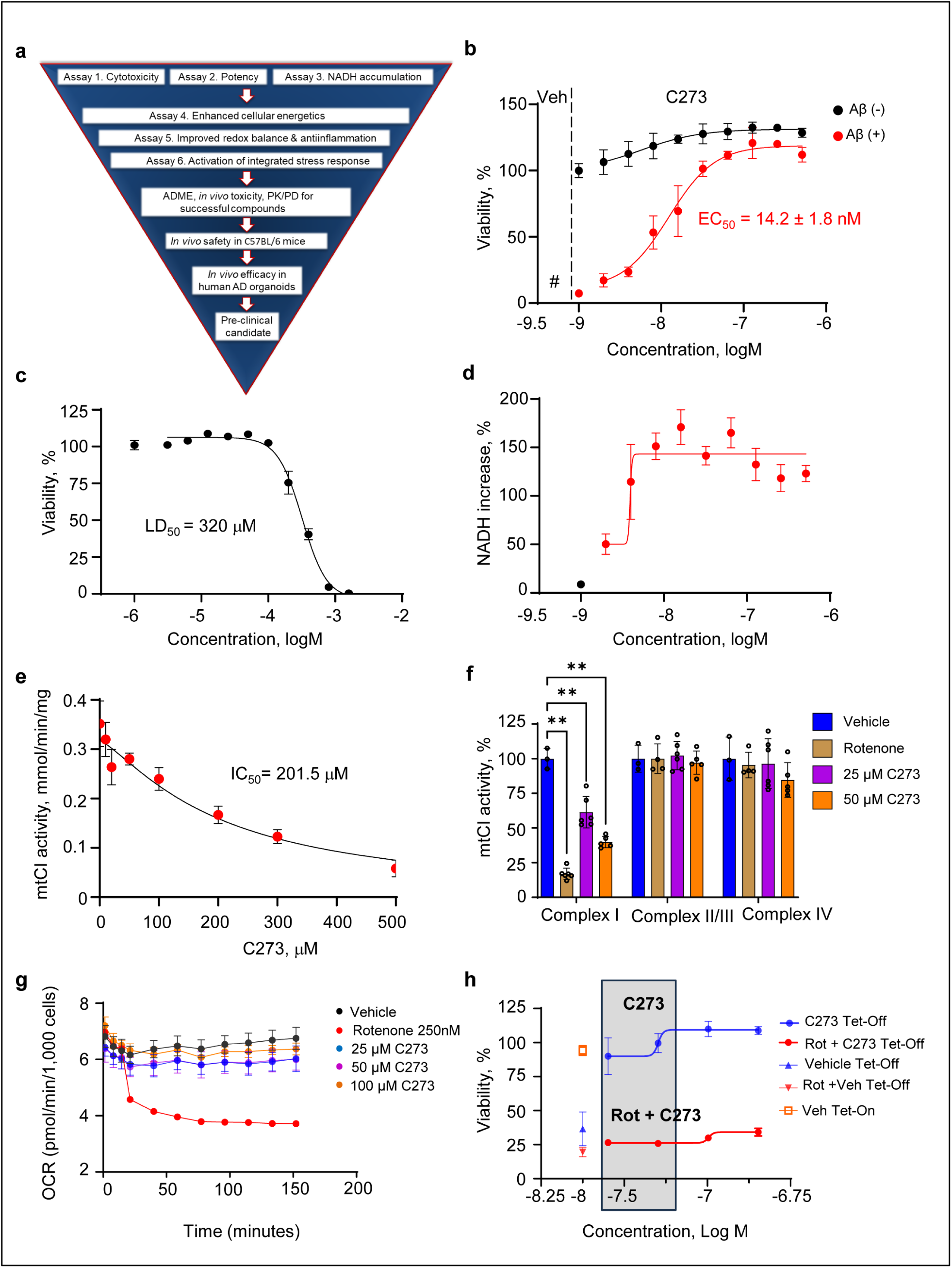
C273 protects against Aβ toxicity through mtCI modulation. **a** A drug discovery funnel used for the identification of C273. **b** C273 protects MC65 Tet-Off cells from Aβ toxicity (red line). # indicates cell death in Tet-Off cells treated with vehicle. No toxicity was observed in MC65 Tet-On cells (Aβ not expressed, black line). EC_50_ values were calculated using nonlinear regression in GraphPad Prism 10. **c** Treatment with C273 for 24 h does not induce cytotoxicity in HepG2 human hepatic cells at concentrations up to 100 μM. **d** C273 increases NADH levels in MC65 Tet-On cells, consistent with inhibition of mtCI. NADH levels were measured over 24 h. **e** Inhibition of mtCI activity in isolated mouse brain mitochondria by C273 measured using an NADH oxidation assay. **f** Activities of mitochondrial ETC complexes I–IV were assessed in permeabilized MC65 cells using an electron flow assay in the Seahorse XFe96 Extracellular Flux Analyzer in the presence of C273 (25 and 50 μM). **g** Injection of rotenone or C273 to monitor OCR kinetics in MC65 Tet-On cells. **h** Pretreatment with rotenone blocks the protective effect of C273 in MC65 Tet-Off cells (Aβ expressed). Open orange squares: Tet-On cells, no Aβ expression, treated with vehicle (positive control, 100% viability). Blue triangles: Tet-Off cells (Aβ expressed), which undergo extensive cell death after 3 days. Tet-Off cells treated with C273 (2 nM - 5 μM; blue line) show complete protection from Aβ toxicity. Solid red triangles: Tet-On cells treated with 32 nM rotenone, which show no toxicity. Open red triangles: Tet-Off cells treated with 32 nM rotenone alone, resulting in complete cell death. Pretreatment with 32 nM rotenone blocks the protective effect of C273 in Tet-Off cells (red line, grey box), supporting a competitive interaction at mtCI. Partial rescue at higher C273 concentrations reflects incomplete occupancy of the mtCI binding site by rotenone, which inhibits ∼50% of mtCI activity at this dose. The grey box indicates the EC_50_ concentration range for C273. *P* values were calculated using unpaired two-tailed Student’s *t*-tests; \*\**P* < 0.01. Data are presented as mean ± SD (*n* = 4 technical replicates). All experiments were replicated at least twice.

### C273 is a selective, weak inhibitor of the mtCI quinone binding site

Target engagement and selectivity of C273 were evaluated by measuring NADH accumulation, mtCI enzymatic activity in isolated mouse brain mitochondria and by assessing mitochondrial respiration in permeabilized MC65 cells using a Seahorse XFe96 Extracellular Flux Analyzer (**Fig. 2d-f; Supplementary Fig. S1a**). Because mtCI inhibition by C273 at efficacious concentrations was modest, intracellular NADH accumulation was used as a sensitive readout of mtCI-dependent NADH oxidation (**Fig. 2d**). C273 induced a concentration-dependent increase in intracellular NADH levels in MC65 Tet-On cells within the same concentration range that protected MC65 Tet-Off cells from Aβ toxicity, supporting functional target engagement (**Fig. 2b, d**). In isolated mouse brain mitochondria and in permeabilized MC65 cells, C273 weakly inhibited mtCI activity with an IC_50_ of 201.5 μM, while showing no detectable inhibition of complexes II, III, or IV, confirming selectivity for mtCI (**Fig. 2e, f**).

Weak mtCI inhibition by C273 was further determined by measuring oxygen consumption rate (OCR) in MC65 Tet-On cells following kinetic injection of C273 or rotenone using a Seahorse XFe96 Analyzer (**Fig. 2g; Supplementary Fig. S1a**). Whereas rotenone rapidly and completely suppressed mitochondrial respiration, C273 caused only a modest reduction in basal OCR. Importantly, no increase in extracellular acidification rate (ECAR) was observed, indicating that C273 did not trigger compensatory activation of glycolysis (**Supplementary Fig. S1a**). Thus, at nanomolar concentrations, C273 weakly inhibits mtCI, protects MC65 cells from Aβ toxicity, and does not induce metabolic stress, closely mimicking the mechanism of action previously described for CP2^28^ and C458^27^.

We previously demonstrated that the compound C458 binds to the quinone (Q) site of mtCI using a competitive inhibition assay with rotenone, a well-characterized Q site inhibitor^27^. To determine whether C273 engages the same binding site, MC65 Tet-Off cells were pretreated with 32 nM rotenone for 30 minutes prior to C273 exposure. This rotenone concentration did not induce cytotoxicity but reduced OCR by approximately 50% within 30 minutes, consistent with partial inhibition of mtCI activity (**Fig. 2g**). Rotenone pretreatment abolished the protective effect of C273 against Aβ-induced toxicity, as evidenced by the loss of C273-mediated EC_50_ activity in MC65 Tet-Off cells (**Fig. 2h**). Rotenone alone did not confer protection against Aβ toxicity at 2-32 nM concentration range, indicating that the observed effect was due to competition at the Q site rather than additive mitochondrial stress. Together, these findings demonstrate that C273 engages the Q site of mtCI and that its weak inhibition of mtCI is required for protection against Aβ toxicity. These data support a mechanism in which selective, reversible modulation of mtCI at the Q site activates a protective mitochondrial stress response while avoiding overt cytotoxicity or metabolic disruption. This separation between potent cellular protection and weak enzymatic inhibition is consistent with a mechanism based on weak targeted modulation of mtCI activity rather than respiratory chain blockade.

### *In vitro* ADME and safety profiling identify C273 as a developable CNS lead

The *in vitro* ADME and safety profile of C273 was evaluated across major human cytochrome P450 (CYP) isoforms, hERG channel inhibition, the CEREP SafetyScreen44 panel, aqueous solubility, plasma protein binding, and microsomal stability (**Fig. 3a-c**). C273 showed minimal inhibition of the major human CYP isoforms CYP1A2, CYP2A6, CYP2B6, CYP2C8, CYP2C9, CYP2C19, CYP2E1, and CYP3A4, with IC_50_ values exceeding 30 μM. Moderate inhibition was observed for CYP2D6 (IC_50_ = 6.51 μM), whereas inhibition of CYP3A4 remained negligible with both midazolam and testosterone probe substrates (**Fig. 3a**). Given the high cellular potency of C273 against Aβ-associated toxicity (EC_50_ = 14.2 nM), this profile met predefined criteria for acceptable CYP liability. Cardiac safety assessment showed low hERG inhibition (IC_50_ = 6.31 μM), within the predefined safety window (**Fig. 3b, Supplementary Fig. S1b**). In the CEREP SafetyScreen44 panel, C273 exhibited a favorable selectivity profile at 10 μM, with detectable off-target interactions limited to a small subset of receptors. These effects occurred at concentrations substantially higher than those required for cellular efficacy, suggesting a low likelihood of off-target-mediated adverse effects at pharmacologically relevant exposures (**Supplementary Table 1**). C273 also exhibited high aqueous solubility (307.02 μM), supporting formulation and exposure. Plasma protein binding studies indicated measurable unbound fractions across species, with recoveries of 47.64% in human plasma, 20.18% in rat plasma, and 13.91% in mouse plasma, indicating species-dependent binding while maintaining free drug levels. C273 also showed species-dependent microsomal stability, with moderate clearance in rat liver microsomes and greater stability in human and mouse microsomes, supporting translational development (**Fig. 3c**). To further address potential concerns with lactic acidosis due to treatment, we measured lactate levels in HepG2 cells treated with C273 for 48 hrs (**Supplementary Fig. S1c**). Compared to a short-term treatment (**Supplementary Fig. S1a**), a longer C273 treatment resulted in a dose-dependent accumulation of extracellular lactate at concentrations above 10 µM, well above the EC_50_ range. Together, these data support C273 as a developable CNS lead with favorable *in vitro* ADME and safety characteristics, including low CYP liability, limited hERG activity, high solubility, and acceptable plasma protein binding. These properties distinguish C273 from earlier compounds CP2 and C458, and support its progression beyond proof-of-concept mitochondrial modulation.

**Fig. 3:**
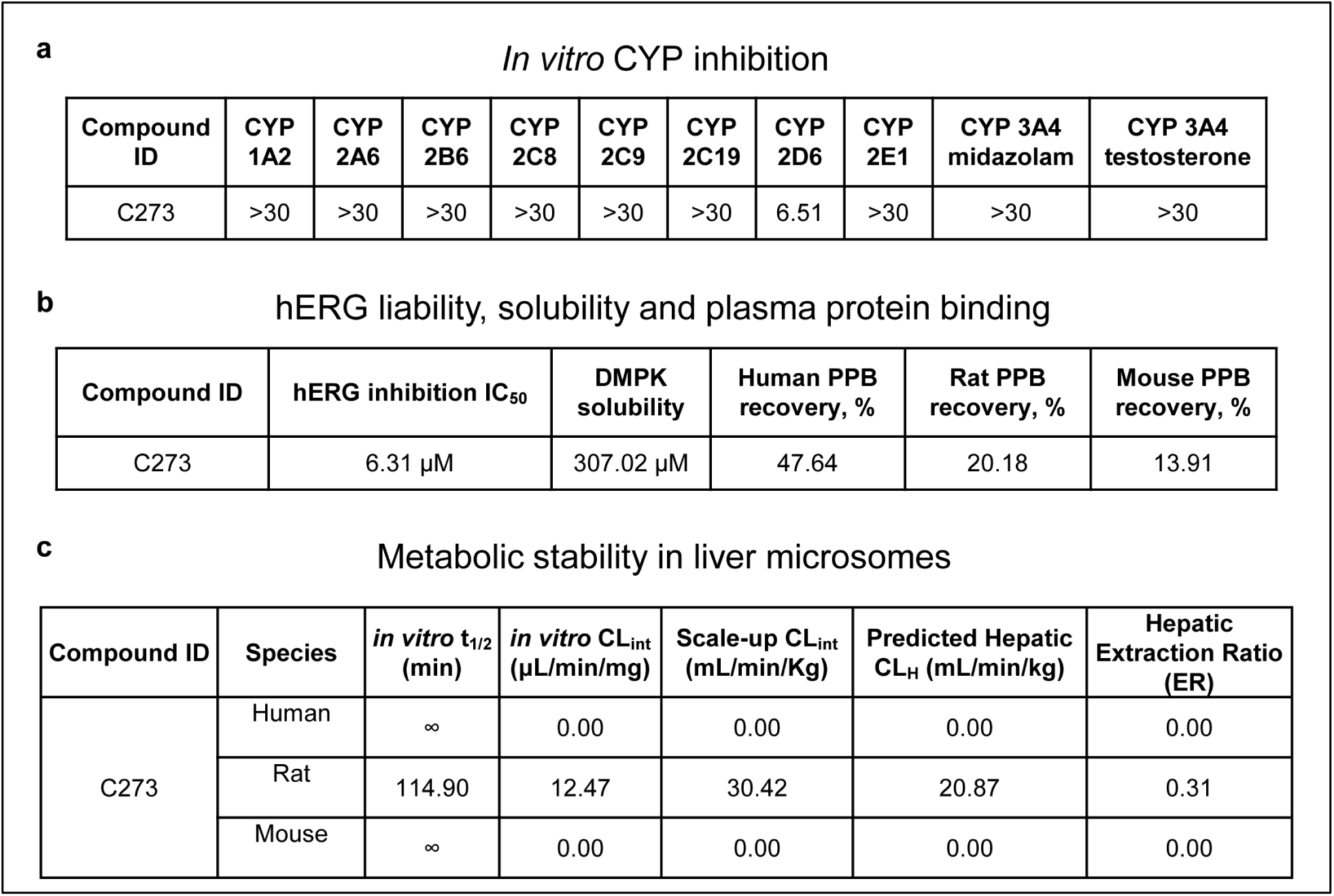
*In vitro* ADME and safety profiling demonstrates favorable developability of C273. **a** Summary of C273 inhibition across major cytochrome P450 (CYP) enzymes. **b** Assessment of hERG channel inhibition, aqueous solubility, and plasma protein binding. **c** Metabolic stability of C273 in liver microsomes.

### C273 shows favorable PK properties, oral bioavailability, and brain penetration *in vivo*

The PK profile of C273 was evaluated in male wild-type (WT) C57BL/6 mice following a single 3 mg/kg intravenous (IV) or 25 mg/kg oral (PO) dose (**Fig. 4a-c**). Plasma and brain samples were collected over 24 h, and C273 concentrations were quantified by LC-MS/MS. C273 demonstrated high brain exposure, with brain-to-plasma ratios of 1.01 after IV dosing and 0.76 after PO dosing. Oral bioavailability was approximately 100% (**Fig. 4c**), indicating efficient systemic absorption and favorable plasma-brain distribution consistent with CNS drug development. After PO administration, C273 exhibited a plasma half-life (T_1/2_) of 1.21 h and a brain half-life of 1.14 h, reaching a maximum plasma concentration (C_max_) of 3615 ng/mL within 30 min. Brain penetration was further supported by a low efflux ratio (ER = 1.65) in MDR1-expressing MDCKII cells, indicating minimal liability for active efflux (**Fig. 4d**). Together, these data show that C273 combines high oral bioavailability with efficient BBB penetration, supporting the feasibility of systemic dosing to achieve therapeutically relevant CNS exposure. The combination of near-complete oral bioavailability and brain-to-plasma ratios approaching unity is notable for a mitochondria-targeted CNS lead compound.

**Fig. 4:**
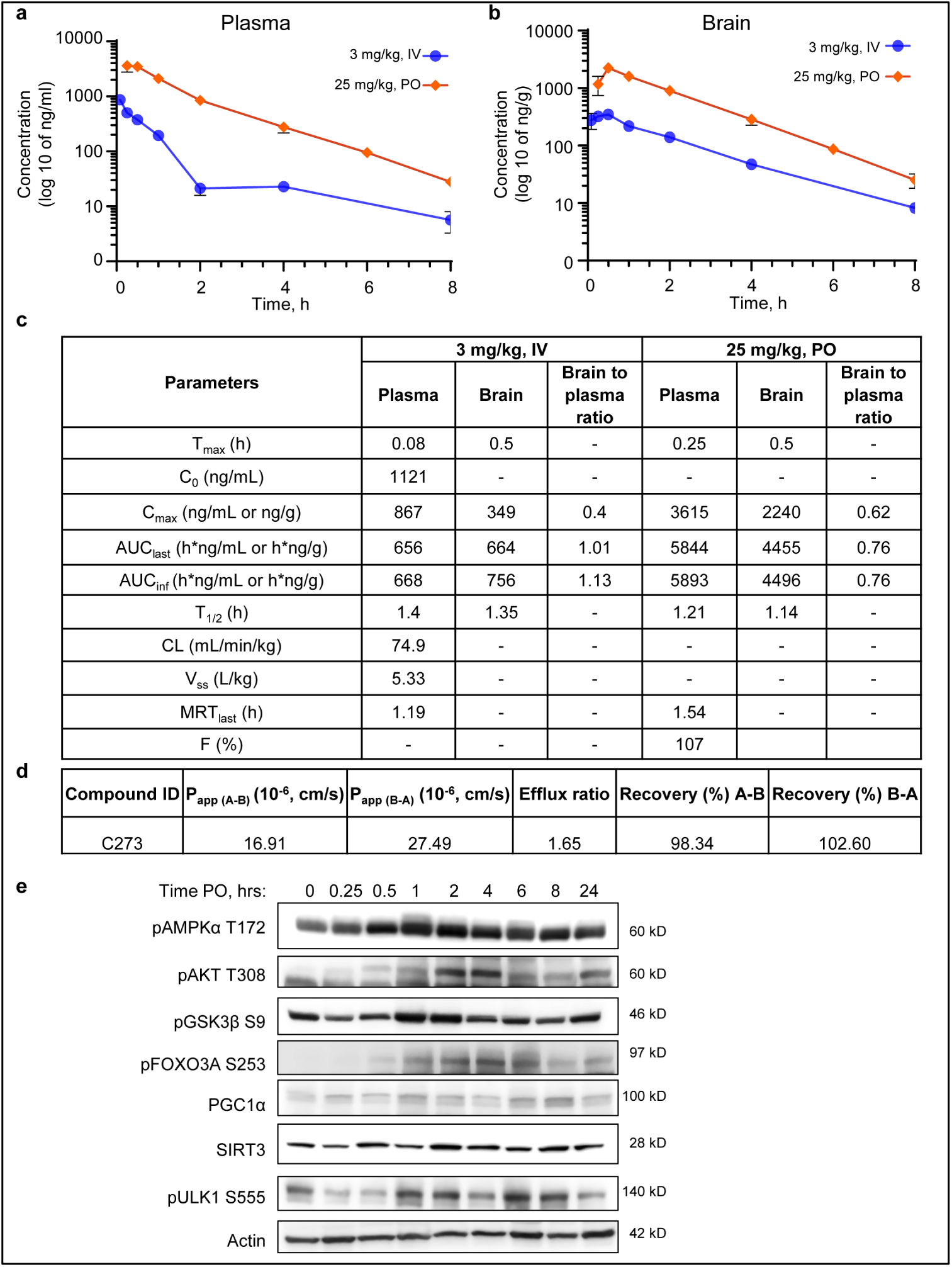
C273 exhibits favorable PK, oral bioavailability, and BBB penetration. **a, b** Pharmacokinetics of C273 in plasma (**a**) and brain (**b**) following intravenous (IV, 3mg/kg) and oral (PO, 25 mg/kg) administration in C57BL/6 male mice (*n* = 3 *per* time point). **c** Summary of PK parameters derived from the studies shown in **a** and **b,** including time to maximum concentration (T_max_), maximum concentration (C_0_ and C_max_), area under the concentration–time curve to the last measurable concentration (AUC_last_), area under the concentration–time curve extrapolated to infinity (AUC_inf_), terminal half-life (t_½_), clearance (CL), elimination rate constant, volume of distribution (V_ss_), mean residence time (MRT), and oral bioavailability (F). **d** Summary of BBB permeability assessed using the MDCK-MDR1 cell assay. **e** Activation of neuroprotective signaling pathways in the mouse brain following a single oral dose of C273 (25 mg/ kg). Western blot analysis was performed on brain tissues collected at multiple time points corresponding to the PK study shown in **b**. Each lane represents an individual mouse.

### C273 activates AMPK-centered neuroprotective signaling in the brain

To assess downstream pharmacology following systemic exposure, Western blot analysis was performed using brain tissue collected during the PK study. Within 30 min of oral administration, C273 increased phosphorylation of AMPKα, indicating rapid activation of a central regulator of cellular energy homeostasis (**Fig. 4e**). Consistent with AMPK activation, phosphorylation of ULK1 at Ser555 was increased, supporting induction of autophagy. C273 also increased Akt phosphorylation, with corresponding inhibitory phosphorylation of downstream targets GSK3β and FOXO3A, pathways implicated in glucose metabolism, tau phosphorylation, and neuronal survival (**Fig. 4e**). In addition, C273 increased brain expression of PGC1α and Sirt3, consistent with stimulation of mitochondrial biogenesis and mitochondrial stress adaptation (**Fig. 4e; Supplementary Fig. S2, S3**). Notably, these signaling effects persisted for up to 24 h after a single oral dose. Together, these results demonstrate that C273 penetrates the BBB and, consistent with CP2^26^ and C458^27^, activates an AMPK-centered neuroprotective program associated with autophagy, mitochondrial biogenesis, and cell survival. These *in vivo* signaling changes support a pharmacological mechanism whereby weak mtCI modulation drives sustained adaptive stress responses in the brain.

### Neuroprotective mechanisms of C273 require AMPK

To determine whether AMPK is required for the protective effects of C273, we used AMPKα1/α2 double-knockout mouse embryonic fibroblasts (MEFs). In WT MEFs, C273 increased phosphorylation of AMPK and its downstream target acetyl-CoA carboxylase (ACC), confirming functional kinase activation (**Fig. 5a**). This was accompanied by increased expression of thioredoxin and IκBα, consistent with enhanced antioxidant defense and suppression of NF-κB signaling (**Fig. 5a**). C273 treatment in WT MEFs also increased expression of PGC-1α, Sirt3, and OXPHOS complexes I and II, consistent with induction of mitochondrial biogenesis (**Fig. 5a; Supplementary Fig. S4, S5)**. In contrast, vehicle-treated AMPKα1/α2-deficient MEFs showed reduced basal levels of PGC-1α, OXPHOS complexes I and V, and phosphorylated ULK1, indicating impaired mitochondrial biogenesis and autophagy. Importantly, C273 failed to induce mitochondrial biogenesis, antioxidant defense, or suppression of inflammatory signaling in AMPKα1/α2-deficient MEFs, demonstrating that AMPK is required for engagement of C273-dependent protective mechanisms. Consistent with these findings, C273 also improved mitochondrial bioenergetic function in SH-SY5Y cells expressing mutant APP (APPswe), significantly increasing spare respiratory capacity, a key indicator of mitochondrial fitness and stress adaptability (**Fig. 5b, c**)^39^. Together, these findings establish AMPK as a key mediator of the neuroprotective effects of C273 and its ability to enhance mitochondrial quality control and bioenergetic function. The genetic requirement for AMPK distinguishes downstream adaptive signaling from nonspecific metabolic suppression and provides mechanistic support for C273 as a signaling-directed mtCI modulator.

**Fig. 5:**
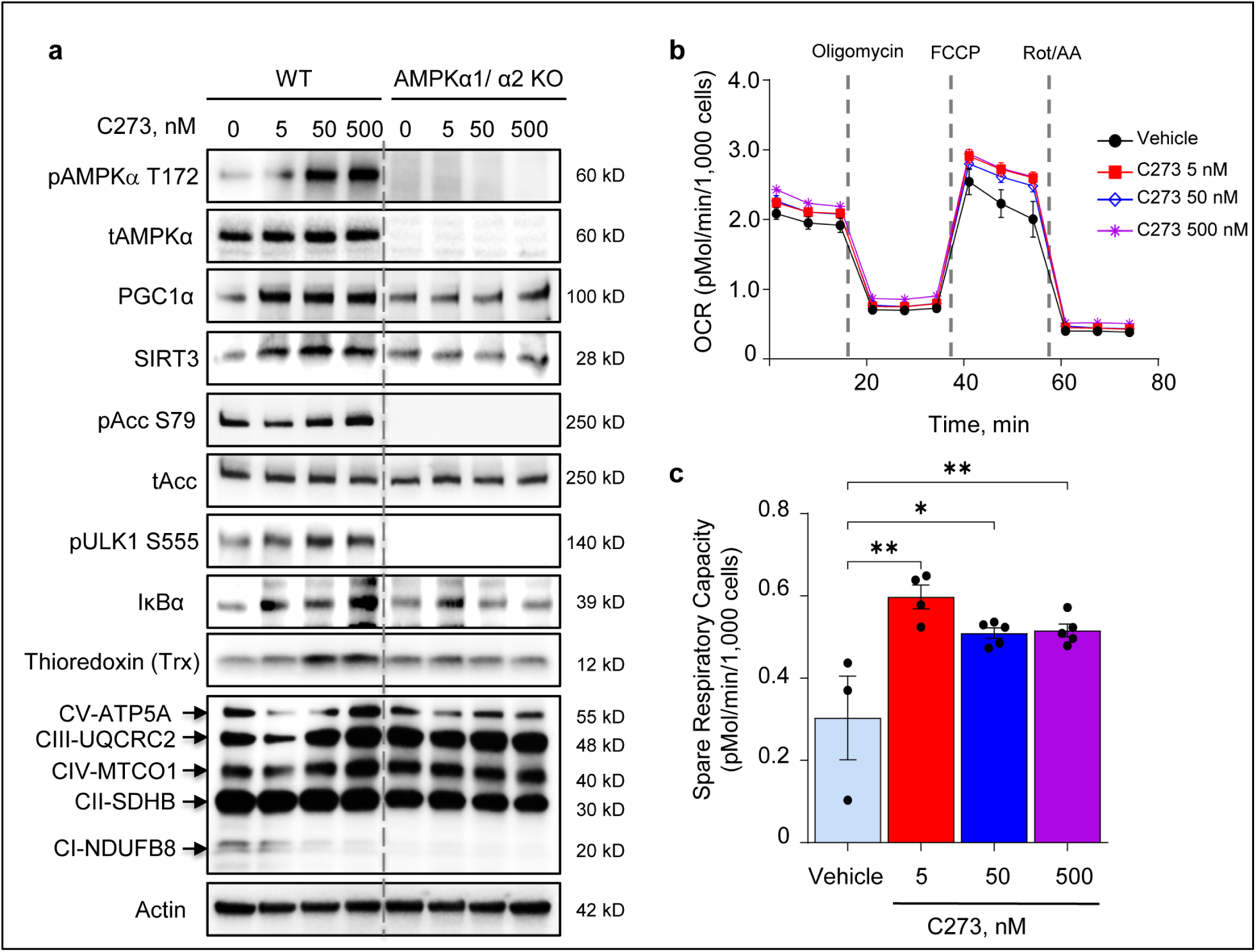
C273 activates AMPK-dependent mitochondrial stress signaling and neuroprotective pathways. **a** C273 treatment activates AMPKα, PGC1α, Sirt3 and autophagy, increases the levels of OXPHOS complexes and antioxidants and attenuates NF-κB signaling in WT MEFs, but not in AMPKα1/α2 KO MEFs. **b** C273 treatment weakly reduces basal OCR and increases SRC in SH-SY5Y APP_SWE_ cells. **c** SRC was calculated from the OCR data shown in **b**. Statistical analysis was performed using two-way ANOVA with Dunnett’s multiple-comparisons test. \**P* < 0.05, \*\**P* < 0.01.

### C273 mitigates oxidative stress and inflammatory signaling

Because AD is associated with elevated oxidative stress and disrupted redox homeostasis, we next examined whether C273 enhances antioxidant defense. In MC65 Tet-Off cells (Aβ is expressed), pretreatment with C273 (250 nM, 24 h) significantly increased resistance to H_2_O_2_-induced toxicity, resulting in an approximately threefold increase in survival (**Fig. 6a**). Importantly, C273 showed no intrinsic antioxidant activity in a cell-free total antioxidant capacity assay (**Supplementary Fig. S6a)**, indicating that its protective effects are not attributable to direct radical scavenging. Consistent with this, C273 increased intracellular NADPH and glutathione (GSH) levels in a dose-dependent manner (**Fig. 6b, c**), reflecting restoration of cellular antioxidant capacity.

**Fig. 6:**
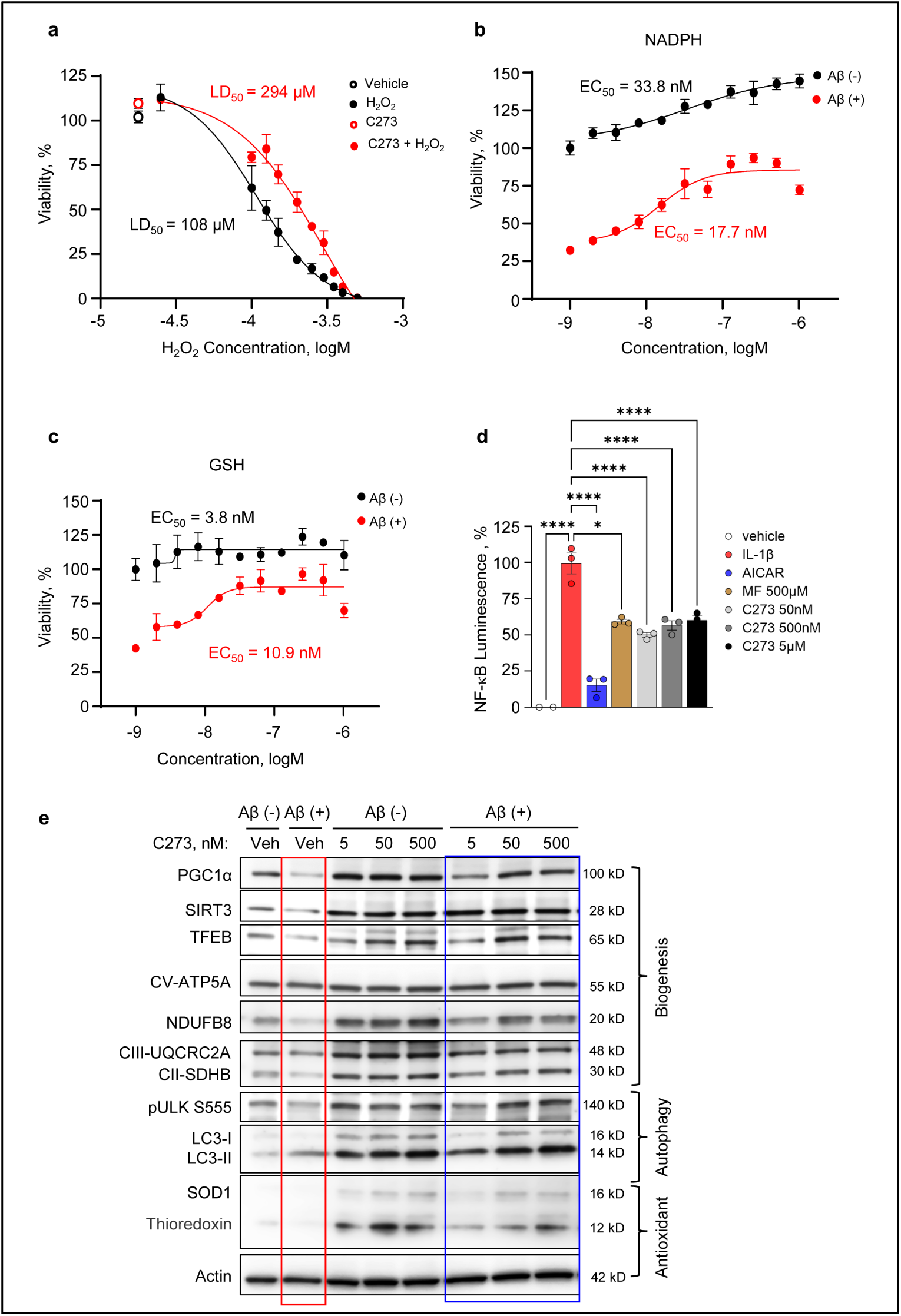
C273 enhances oxidative stress resistance, restores redox balance, and promotes mitochondrial quality control. **a** C273 increases resistance to oxidative stress induced by H_2_O_2_ in MC65 Tet-Off cells expressing Aβ. Cells were pretreated with C273 prior to H_2_O_2_ exposure, and cell viability was assessed to determine LD_50_ values. LD_50_ values were calculated by nonlinear regression using GraphPad Prism. **b, c** C273 restores intracellular NADPH (**b**) and GSH (**c**) levels in MC65 Tet-Off cells. **d** C273 suppresses IL-1β-induced NF-κB activation. NF-κB reporter HEK293 cells were pretreated for 24 h with C273 (50 nM, 500 nM, or 5 μM), AICAR (0.5 mM), or metformin (0.5 mM), followed by stimulation with IL-1β (1 ng/ml) for 6 h. NF-κB luciferase activity was measured using the Luciferase Assay System (Promega, E1501). *n* = 4 independent experiments. **e** C273 restores mitochondrial biogenesis, expression of ETC complexes, autophagy markers, and antioxidant proteins in MC65 Tet-Off cells. MC65 cells were cultured under Tet-On (Aβ-) or Tet-Off (Aβ+) conditions and treated with vehicle or C273 (5 nM, 50 nM, or 500 nM) for 24 h. Protein levels were analyzed by Western blot. Statistical analysis was performed via one-way ANOVA to compare the vehicle-, metformin-, AICAR- and C273-treated groups. Data are presented as mean ± SD. \**P* < 0.05; \*\**P* < 0.01.

We next assessed inflammatory signaling using HEK293 NF-κB reporter cells stimulated with IL-1β or TNFα. C273 markedly suppressed NF-κB activation, with anti-inflammatory activity comparable to that of metformin and AICAR, two established AMPK activators (**Fig. 6d; Supplementary Fig. S6b**). These data indicate that C273 suppresses pro-inflammatory signaling in parallel with the restoration of antioxidant defenses.

Intracellular Aβ accumulation in MC65 Tet-Off cells impaired the expression of proteins involved in mitochondrial biogenesis, autophagy, and antioxidant defense, including TFEB, PGC1α, Sirt3, OXPHOS complexes, as well as decreased phosphorylation of ULK1, reduced LC3-II levels, and diminished expression of superoxide dismutase (SOD) and thioredoxin (**Fig. 6e**). Treatment of MC65 Tet-Off cells with C273 restored the expression of these key regulatory proteins and rescued cells from Aβ-mediated cytotoxicity (**Fig. 6e; Supplementary Fig. S7, S8**). Together, these data show that weak mtCI inhibition by C273 promotes antioxidant defense, suppresses inflammatory signaling, and restores mitochondrial quality control pathways. These coordinated effects are consistent with pharmacological engagement of an adaptive mitochondrial stress response rather than direct antioxidant activity alone.

### Preliminary toxicology of C273 in WT mice

To assess *in vivo* tolerability, 2-month-old female WT mice were treated with vehicle or C273 at 20, 50, or 80 mg/kg/day in drinking water for 30 days (*n* = 5 *per* group). Body weight gain was comparable across all groups throughout the treatment period (**Fig. 7a**), and locomotor activity in the open field test was unchanged (**Fig. 7b**), indicating preserved general health and behavior. After one month of treatment, mice were sacrificed, and major organs were collected. Liver and heart were examined histologically. No tissue pathology, inflammatory changes, or structural abnormalities were detected in C273-treated mice at any tested dose (**Fig. 7c; Supplementary Fig. S9**). No systemic adverse effects were observed. These results indicate that C273 is well tolerated *in vivo* and does not produce detectable cardiac or hepatic toxicity following 30 days of administration at doses up to 80 mg/kg/day. This tolerability window is consistent with the weak and controlled mode of mtCI inhibition achieved by C273.

**Fig. 7:**
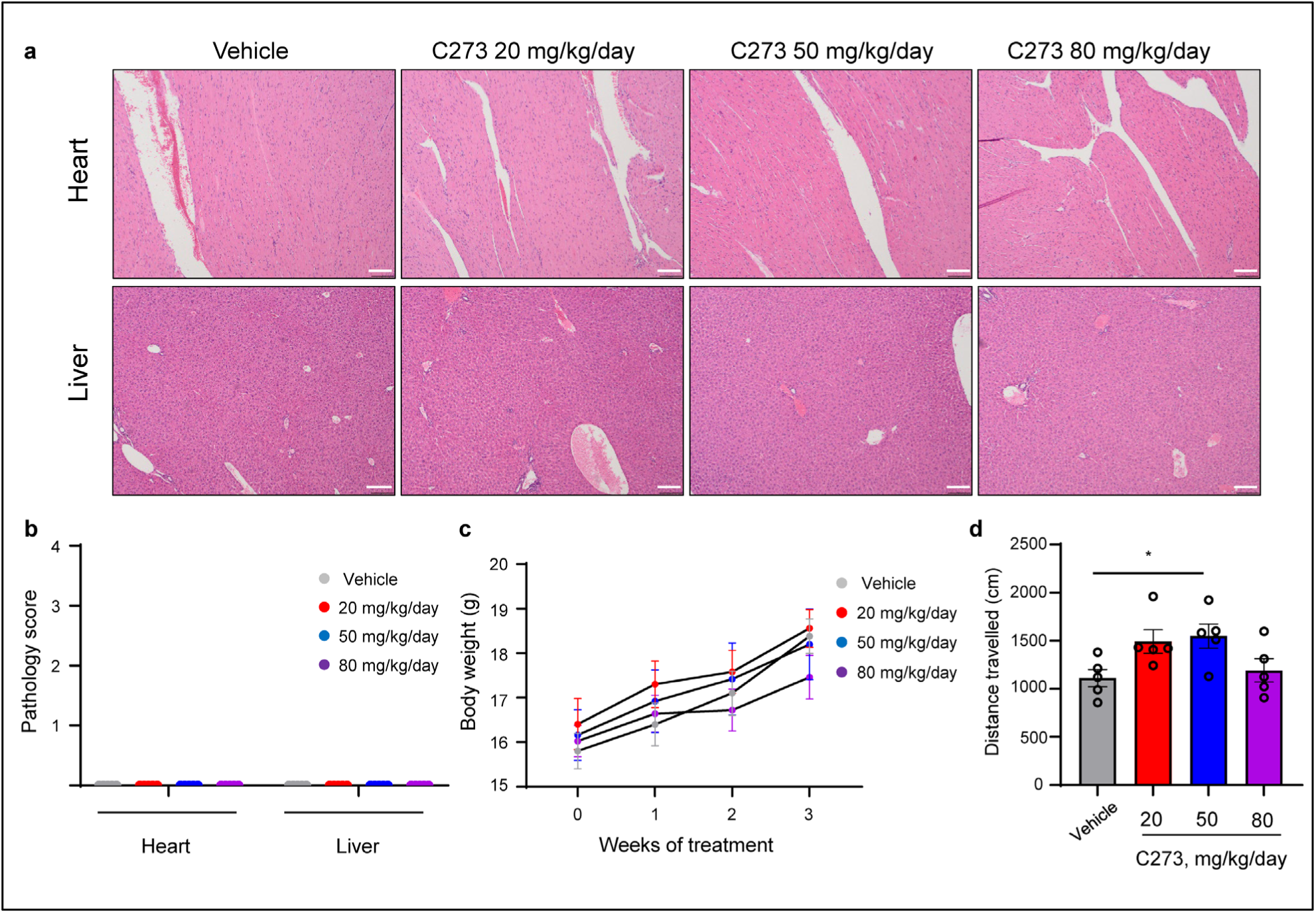
C273 shows no detectable toxicity in WT mice. **a** Representative histology sections of heart and liver tissues from WT mice treated with different doses of C273 stained with H&E, showing no detectable pathological changes. Scale bars, 500 µm. **b** Pathology scores determined by an independent pathologist blinded to treatment groups demonstrated the lack of pathological changes due to treatment. **c** Weekly body weight measurements during the treatment period. **d** Locomotor activity assessed using the Open Field Test. *N* = 5 mice *per* group. Treatment was conducted for 30 days. Statistics was done using the ordinary one-way ANOVA with the Dunnett-corrected post-hoc for the multiple comparisons. \**P* < 0.05.

### Peripheral AMPK activation as a translational pharmacodynamic biomarker

To establish a translational pharmacodynamic biomarker of mtCI engagement, we assessed AMPK activation in peripheral blood cells following exposure to C273 and CP2. Because AMPK activation is the proximal signaling event linking mtCI modulation to downstream mitochondrial remodeling and antioxidant defense, we measured pAMPK Thr172 together with PGC1α, OXPHOS complexes, and antioxidant enzymes as a coordinated pharmacodynamic signature (**Fig. 8a-c**). Western blot analyses were performed using mouse peripheral blood mononuclear cells (PBMCs) isolated from CP2-gavaged mice and human PBMCs treated with CP2 *in vitro* (**Fig. 8a, b; Supplementary Fig. S10**, **S11)**. CP2 induced a similar response, activating AMPK and the downstream signaling cascade, leading to increased expression of PGC1α and antioxidants, including SOD1 and thioredoxin (**Fig. 8a, b**). This data supports the translational relevance of the mouse data.

**Fig. 8:**
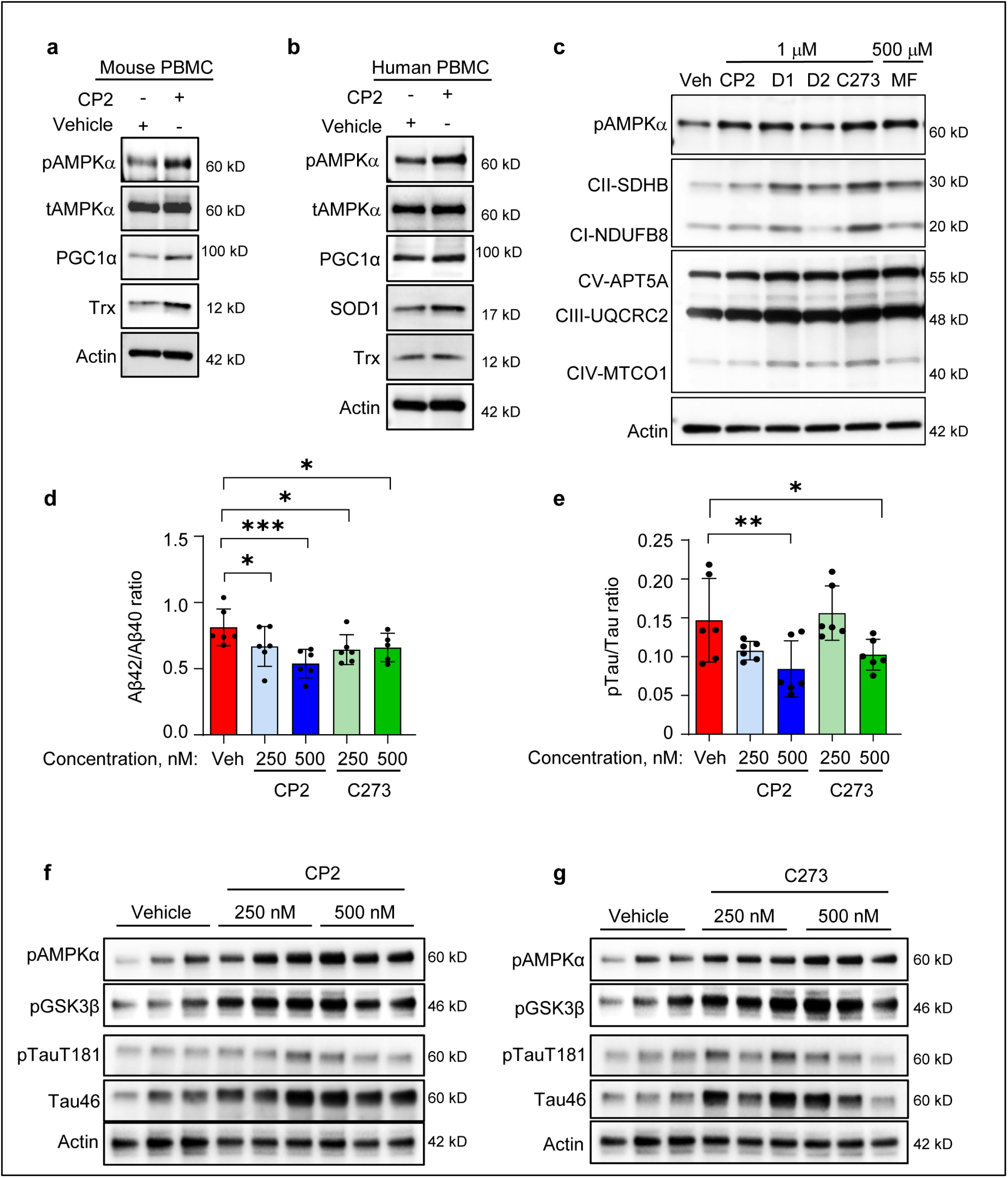
mtCI inhibitors activate target engagement biomarkers in PBMCs and reduce Aβ and pTau in human LOAD organoids. **a** PBMCs were isolated from three female WT mice treated by oral gavage with CP2 (25 mg/ kg) or vehicle (20% PEG400 in D5W). Blood was collected 24 h post-treatment, pooled, PBMCs were isolated by Ficoll density-gradient centrifugation, lysed, and protein extracts were analyzed by Western blot. **b,c** Human PBMCs were isolated from the whole blood of a healthy donor. Cells were treated for 24 h with vehicle (0.01% DMSO) or mtCI inhibitors: CP2, CP2 diastereomer D1, CP2 diastereomer D2, C273 (all at 1 μM), or metformin (MF, 500 μM). Protein levels were assessed by Western blot. **d-e** The levels of Aβ40, Aβ42, p-Tau (T181), and total Tau in RIPA fractions from human iPSC-derived organoids treated with vehicle, CP2 or C273 were measured by ELISA. The data were normalized to the total protein concentration of each sample (*n* = 3 organoids *per* treatment, measured in duplicate). Data are expressed as mean ± SD. Statistical analysis was performed using one-way ANOVA with the FDR-corrected post hoc for multiple comparisons. \**P* < 0.05, \*\**P* < 0.01. **f,g** Western blots of human iPSC-derived organoids treated with vehicle or different doses of CP2 and C273.

Treatment of human PBMCs with C273, CP2, the CP2 diastereoisomers D1 and D2, and metformin induced pronounced phosphorylation of pAMPK at Thr172 (**Fig. 8c; Supplementary Fig. S12**). Notably, at 1 μM C273 induced AMPK activation comparable to that achieved with metformin at 500 μM, indicating substantially higher potency.

Peripheral AMPK activation by C273 was accompanied by induction of PGC1α, increased OXPHOS complex abundance, and upregulation of antioxidant enzymes, consistent with engagement of an AMPK-driven adaptive mitochondrial stress response. Importantly, peripheral pAMPK Thr172 activation paralleled the increase in brain pAMPK Thr172 observed in C273-treated mice in the present study and in CP2-treated mice reported previously^25–28^, supporting a coordinated central and peripheral pharmacodynamic response to weak mtCI inhibition. Together, these findings support the use of pAMPK Thr172 activation in PBMCs as a minimally invasive translational biomarker of target engagement and downstream pharmacodynamic activity for C273-class compounds. The identification of a peripheral biomarker linked to brain target engagement is an important translational feature for future clinical development.

### C273 and CP2 reduce Aβ and p-Tau levels in patient-derived LOAD cerebral organoids

To assess translational relevance in a human disease context, we generated induced pluripotent stem cell-derived cerebral organoids from patients with late-onset AD (LOAD) carrying the APOE4/4 alleles ^40^. These three-dimensional organoids recapitulate key features of AD-related pathology, including increased levels of Aβ and p-Tau (T181). Organoids were treated with C273 or CP2 for 48 h, followed by quantitative analysis of Aβ40, Aβ42, p-Tau (T181), and total tau by ELISA (**Fig. 8d, e; Supplementary Fig. S13**). C273 significantly reduced the intracellular Aβ42/Aβ40 and the p-Tau (T181)/t-Tau ratios relative to vehicle-treated controls. CP2 produced a similar reduction in both pathological markers, consistent with its established mtCI-mediated neuroprotective mechanism^27^. The reduction in the Aβ42/Aβ40 ratio is particularly notable in the APOE4/4 background, which is associated with impaired amyloid clearance and preferential accumulation of Aβ42^41^. To confirm target engagement and downstream signaling in this human system, Western blot analysis was performed following treatment with C273 or CP2 in two independent experiments in two independent experiments (**Fig. 8f,g; Supplementary Fig. S14, S15, S16**). Both compounds robustly increased pAMPKα at T172 and pGSK3β at S9, indicating activation of AMPK signaling and inhibitory phosphorylation of GSK3β. Consistent with the ELISA results, both C273 and CP2 also reduced Tau phosphorylation at T181 **(Supplementary Fig. S14c)** . Given the established role of GSK3β in tau phosphorylation,^42,43^, these findings provide a mechanistic link between mtCI modulation, AMPK activation, and the observed reduction in levels of p-Tau.

Together, these data demonstrate that weak mtCI inhibition by C273 and CP2 activates neuroprotective signaling pathways and mitigates core molecular hallmarks of AD in human patient-derived cerebral organoids. The convergence of reduced Aβ and p-tau levels with activation of the AMPK/ GSK3β axis supports adaptive mitochondrial modulation as a translationally relevant therapeutic strategy. These findings extend the pharmacology of C273 beyond rodent and immortalized cell systems and establish its activity in a human, genetically high-risk AD background.

## Discussion

Efforts to develop disease-modifying therapies for AD focused on amyloid-centered approaches yielded limited success^44–47^. Increasing evidence indicates that mitochondrial dysfunction, oxidative stress, metabolic failure, and chronic neuroinflammation represent early and persistent drivers of disease progression^3,48^. Therapeutic targeting of mitochondrial metabolism has therefore emerged as a promising alternative strategy^49^. Mitochondrial inhibitors used clinically or experimentally, including classical mtCI inhibitors such as rotenone and oncology-directed compounds such as IACS-010759, produce strong inhibition of the ETC that leads to bioenergetic collapse and toxicity^50,51^. In contrast, recent work suggests that weak inhibition of mtCI can trigger protective adaptive stress responses without compromising cellular energy production. One of the examples includes metformin, an FDA-approved drug for type 2 diabetes, which is a weak but non-specific mtCI inhibitor^52,53^. Previous studies from our group^10,25–32,54^ demonstrated that the small molecule CP2 weakly inhibits mtCI and ameliorates AD pathology and cognitive deficits in multiple AD mouse models, establishing proof of concept that controlled modulation of mitochondrial respiration can confer therapeutic benefit. Notably, CP2 also restored mitochondrial and neuronal network function in a mouse model with ∼50% loss of mtCI activity, the *Ndufs4* knockout mice, supporting both the safety and efficacy of this approach under conditions of severe mitochondrial dysfunction^30^.

In the present study, we advance this strategy by identifying and characterizing C273, a clinically optimized mtCI modulator derived from iterative medicinal chemistry optimization of CP2 and C458^27^. This work defines a coherent drug discovery trajectory in which CP2 established proof of concept for the therapeutic potential of weak mtCI inhibition, C458 improved potency and selectivity, and C273 incorporates chemical and pharmacological properties suitable for translational development while retaining the core mechanistic features of this chemotype. Medicinal chemistry optimization improved drug-like properties while preserving mtCI engagement. Structural modification of the parent scaffold reduced stereochemical complexity and enabled the synthesis of C273 as a single S-enantiomer with improved yield. These changes preserved nanomolar protection against Aβ-induced toxicity in a phenotypic screen while producing favorable physicochemical and ADME characteristics, including improved solubility, acceptable plasma protein binding, minimal CYP and hERG liabilities, and microsomal stability compatible with predicted human pharmacokinetics. Together with robust oral bioavailability and efficient BBB penetration, these properties position C273 as a pharmacologically optimized representative of the mtCI-modulating chemotype.

Mechanistically, C273 acts as a weak and selective inhibitor of mtCI. In contrast to classical inhibitors such as rotenone, which binds tightly to mtCI and completely blocks ETC with rapid bioenergetic collapse, C273 produces only modest inhibition of mtCI activity and a mild reduction in oxygen consumption without inducing metabolic stress or compensatory glycolysis. This distinction is critical because excessive mtCI inhibition is known to be neurotoxic. Instead, weak inhibition appears to relieve pathological electron pressure within the respiratory chain, thereby limiting mitochondrial ROS production while preserving sufficient respiratory capacity to sustain cellular energy demands. Target engagement was supported by concentration-dependent accumulation of NADH and by competitive replacement by rotenone, consistent with modulation of NADH oxidation by mtCI. These findings are supportive of interaction at the Q binding site of mtCI, a critical region where the final steps of electron transfer from NADH to ubiquinone is localized. Engagement of this site enables fine-tuning of electron flux without completely blocking the respiratory chain and likely contributes to C273’s high selectivity for mtCI over other respiratory complexes.

A central mechanistic insight from this work is that the neuroprotective activity of C273 depends on AMPK activation. AMPK plays an important role in AD, and its activation has been shown to reverse multiple pathophysiological pathways^55–57^. Genetic ablation of AMPKα1/α2 abolished C273-induced mitochondrial biogenesis, antioxidant responses, autophagy, and suppression of inflammatory signaling, demonstrating that AMPK is required for the adaptive response triggered by weak mtCI inhibition. In WT cells, C273 activated AMPK and stimulated pathways involved in mitochondrial quality control and metabolic adaptation, including phosphorylation of acetyl-CoA carboxylase, induction of autophagy through ULK1 activation, and mitochondrial biogenesis through PGC-1α and Sirt3. These responses were accompanied by increased expression of OXPHOS complexes and improved mitochondrial bioenergetic capacity. Importantly, mtCI-dependent AMPK activation differs mechanistically from pharmacological AMPK activators such as AICAR or Compound-13^58^, which activate AMPK independently of mitochondrial context. By modulating mitochondrial electron flux, C273 engages a localized mitochondrial stress signaling program that couples AMPK activation to mitochondrial maintenance, redox homeostasis, and metabolic adaptation^14,20–24,59,60^. Consistent with this mechanism, C273 enhanced resistance to oxidative stress and increased intracellular NADPH and glutathione levels in Aβ-stressed neuronal models. Increased expression of endogenous antioxidant enzymes further indicated reinforcement of intrinsic redox defense systems. In parallel, C273 suppressed NF-κB-dependent inflammatory signaling in an AMPK-dependent manner, supporting a model in which correction of mitochondrial dysfunction attenuates inflammatory pathways that contribute to neurodegeneration. The indirect activation of AMPK by mtCI modulators distinguishes this therapeutic approach from direct activators, which have encountered multiple problems in clinical translation^61^. This unique approach also may eliminate controversies associated with the AMPK activation in AD^62^.

An important translational aspect of this work is the identification of peripheral AMPK activation as a pharmacodynamic biomarker of mtCI engagement. Both CP2 and C273 induced AMPK phosphorylation in mouse and human PBMCs, and oral CP2 administration increased p-AMPK levels in peripheral blood *in vivo*. Peripheral activation paralleled AMPK activation in brain tissue, suggesting that PBMC-based monitoring of AMPK phosphorylation may serve as a minimally invasive biomarker of target engagement for clinical development. Together with prior studies demonstrating that CP2 efficacy can be monitored *in vivo* in APP/PS1 mice using FDG-PET^26^, these findings support a set of translational biomarkers for both target engagement and therapeutic efficacy.

Pharmacokinetic studies demonstrated that C273 possesses properties favorable for CNS drug development, including excellent oral bioavailability and efficient BBB penetration with brain-to-plasma ratios close to unity. Despite a relatively short plasma half-life, C273 induced sustained activation of AMPK and mitochondrial adaptive stress pathways in brain tissue, consistent with a pharmacodynamic mechanism driven by adaptive signaling rather than continuous target occupancy. This profile suggests that intermittent dosing may achieve durable therapeutic effects while minimizing safety concerns associated with chronic mitochondrial inhibition. The therapeutic relevance of mtCI modulation was further supported in human iPSC-derived cerebral organoids generated from AD patients carrying APOE4/4 alleles. In this model, both C273 and CP2, similar to C458, reduced the Aβ42/Aβ40 and the pTau/Tau ratios, extending previous observations in transgenic mouse models and demonstrating that weak mtCI inhibition mitigates key molecular hallmarks of AD in human neural tissue^27^. Importantly, both compounds increased phosphorylation of AMPK at T172 and GSK3β at S9, confirming target engagement and activation of a key neuroprotective signaling axis in this human system. Given the central role of GSK3β in tau phosphorylation, its inhibitory phosphorylation provides a direct mechanistic link between mtCI modulation, AMPK activation, and the observed reduction in p-Tau levels, as we observed in Aβ and pTau mouse models^25–27^. Collectively, these results establish C273 as a clinically optimized member of a new class of mtCI modulators that induce protective metabolic stress signaling rather than direct targeting of individual pathogenic pathways. By selectively engaging the Q binding site of mtCI, weak inhibition of electron flux produces a mild bioenergetic shift that activates AMPK-dependent adaptive stress signaling, leading to restoration of mitochondrial function, improved redox balance, enhanced autophagy, and suppression of inflammatory pathways. This mechanism links mitochondrial target engagement to downstream correction of amyloid and tau pathology, supporting mtCI modulation as a viable disease-modifying strategy for AD.

## Methods

### Ethics declaration

All experiments with mice were approved by the Mayo Clinic Institutional Animal Care and Use Committee in accordance with the National Institutes of Health’s *Guide for the Care and Use of Laboratory Animals* and ARRIVE guidelines. IACUC protocol number A00001186.

### Reagents

CP2 was synthesized by Nanosyn, Inc. (http://www.nanosyn.com), as described previously, ^63^ and was purified using HPLC. Authentication was performed through NMR spectra to ensure the lack of batch-to-batch variation in purity. CP2 was synthesized as a free base. CP2 synthesis yields a racemic mixture of diastereoisomers, which were purified using chiral chromatography by CRO Charnwood Molecular (UK). For *in vitro* experiments, CP2 was prepared as a 10 mM stock solution in DMSO. Stock aliquots of 20 µl were stored at -80 °C. The following reagents were used: DMSO (Sigma, D2650), rotenone (Sigma, R8875), metformin (Cayman Chemical, 13118), hydrogen peroxide 30% solution (Sigma, H1009), MEM nonessential amino acids 100x (Corning, 25-025-CI), penicillin‒streptomycin (Sigma, PO781), tetracycline hydrochloride (Sigma, T7660), high-glucose DMEM (Thermo Scientific, 11995065), heat-inactivated fetal bovine serum (Sigma, F4135), RPMI-1640 (Corning, MT10041CM), DPBS 1x (Corning, 21-031-CM), sodium pyruvate (Corning, MT25000CI), and phenol red-free Opti-MEM^TM^ I Reduced Serum Medium (Thermo Scientific, 51200038).

### Antibodies

The following primary antibodies were used: p-AMPKα (Thr 172) (1:1000, Cell Signaling Technology, cat. # 2535, RRID:AB_331250), AMPKα (1:1000, Cell Signaling Technology, cat. # 2532, RRID:AB_330331), p-acetyl-CoA carboxylase (Ser79) (1:1000, Cell Signaling Technology, cat. #11818, RRID:AB_2687505), p-GSK3β (Ser 9) (1:1000, Cell Signaling Technology, cat. # 9323, RRID:AB_2115201), GSK3β (1:1000, Cell Signaling Technology, cat. # 9832, RRID:AB_10839406), Sirt 1 (D1D7) (1:1000, Cell Signaling Technology, cat. # 9475S), Sirt3 (1:1000, Cell Signaling Technology, cat. # 5490, RRID:AB_10828246), Superoxide Dismutase 1 (1:1000, Abcam, cat. # ab16831, RRID:AB_302535), p-Akt (Ser473) (1:1000, Cell Signaling Technology, cat. # 4051, RRID:AB_331158), and p-Akt (Thr308) (1:1000, Cell Signaling Technology, cat. # 4056, RRID:AB_331163), p-FOXO1A (Cell Signaling Technology, cat. # 84192, RRID:AB_2800035), p-FOXO3A (Cell Signaling Technology, cat. # 9466, RRID:AB_2106674), p-ULK1 (Ser555) (1:1000, Cell Signaling Technology, cat. # 5869, RRID:AB_10707365), p-ULK1 (Ser317) (1:1000, Cell Signaling Technology, cat. # 12753, RRID:AB_2687883), PGC1α (4C1.3) ST1202 (1∶1000, Calbiochem, cat. # KP9803), PGC-1α (3G6) (1:1000, Cell Signaling Technology, cat. #2178), OXPHOS cocktail (Abcam, ab110413, RRID:AB_2629281), NF-κB Pathway Sampler Kit (Cell Signaling Technology, 9936T, RRID:AB_561197), IκBα (1:1000, Cell Signaling Technology, cat. # 4812, RRID:AB_10694416), HO-1 (1:1000, Cell Signaling Technology, cat. # 70081, RRID:AB_2799772), Oxidative Stress Defense Cocktail (1:250, Abcam, cat. #ab179843), RRID:AB_628036NDUFA9 (Thermo Fisher Scientific Cat# 459100, RRID:AB_10376187), p-Tau T181 (1:200, AT270, Thermo Fisher/Invitrogen MN1050), RRID:AB_223651, Total Tau46 (Cell Signaling #4019), RRID:AB_10695394, Anti-β-Actin (1:5000, Sigma-Aldrich, cat. # A5316, RRID:AB_476743). The following secondary antibodies were used: donkey anti-rabbit IgG conjugated with Horseradish Peroxidase (1:10000 dilution, GE Healthcare UK Limited, UK) and sheep anti-mouse IgG conjugated with horseradish peroxidase (1:10000 dilution; GE Healthcare UK Limited, UK).

### Cells

Human neuroblastoma MC65 cells were a gift from Dr. Bryce Sopher (University of Washington, Seattle, USA). Expression of C99 fragment in MC65 cells was confirmed by Western blot following by cell death measurements at 72 hours at Tet-Off (Ab is expressed) conditions. AMPKα1/a2 knockout mouse embryonic fibroblasts (MEFs) were a gift from Dr. Benoit Viollet (Inserm, Paris, France). The lack of AMPKα1/α2 expression was confirmed by Western blot. Human neuroblastoma SH-SY5Y cells stably transfected with Swedish mutant amyloid precursor protein (APP_SWE_) were a gift from Dr. Cristina Parrado (Karolinska Institute, Sweden). Authentication of SH-SY5Y APP_SWE_ cells was confirmed by real time PCR. The cells were grown in high-glucose DMEM supplemented with 10% FBS, 1 mM sodium pyruvate and 1x nonessential amino acids. NF-κB Reporter (Luc) HEK293 cell line and the antioxidant response element (ARE) Luciferase Reporter HepG2 hepatic cell line were purchased from BPS Bioscience (CA, USA) and were cultured in accordance of manufacturer instructions. All cells were incubated in 5% CO_2_ at 37 °C unless otherwise noted. Cells were regularly checked and were confirmed to be negative for mycoplasma contamination.

### Isolation of peripheral blood mononuclear cells from mouse and human samples

Peripheral blood mononuclear cells (PBMCs) were isolated from three female wild-type mice treated by oral gavage with CP2 (25 mg/kg) or vehicle (20% PEG400 in D5W). Blood was collected 24 h post-treatment, pooled, and PBMCs were isolated by Ficoll density-gradient centrifugation.

Human PBMCs were obtained from apheresis cones of healthy donors through the Division of Transfusion Medicine at Mayo Clinic (Rochester, Minnesota) in accordance with current regulations of the AABB and the U.S. Food and Drug Administration. PBMCs were isolated by Ficoll density-gradient centrifugation as previously described ^64^.

### Synthesis of a mtCI inhibitor C273

#### Preparation of *tert*-Butyl (*S*)-(1-Oxo-1-(4-(*m*-tolyloxy)piperidin-1-yl)propan-2-yl)carbamate

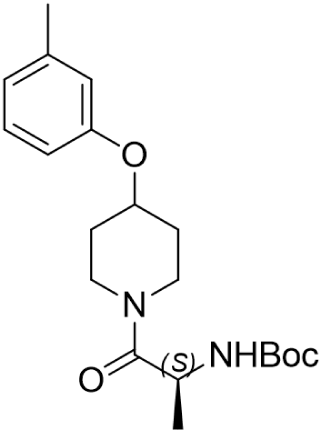

To a stirred room temperature solution of 4-(*m*-tolyloxy)piperidine (6.72 g, 35.1 mmol) in dichloromethane (167 mL) under a nitrogen atmosphere was added *N*-ethyl-*N*-isopropylpropan-2-amine (12.2 mL, 70.3 mmol) and (*tert*-butoxycarbonyl)-*L*-alanine (7.98 g, 42.2 mmol). To the resulting solution was added 2,4,6-tripropyl-1,3,5,2,4,6-trioxatriphosphinane-2,4,6-trioxide (31.1 mL, 52.7 mmol) and the reaction mixture stirred at room temperature for 16 h. After this time, the reaction mixture was concentrated onto silica gel, and purified by column chromatography (120 g, silica, 0 to 30% ethyl acetate/dichloromethane) to afford *tert*-butyl (*S*)-(1-oxo-1-(4-(*m*-tolyloxy)piperidin-1-yl)propan-2-yl)carbamate (11.1 g, 87%) as a clear oil: ¹H NMR (300 MHz, CDCl3) δ 7.17 (t, *J* = 7.7 Hz, 1H), 6.93–6.66 (m, 3H), 5.57 (d, *J* = 7.8 Hz, 1H), 4.71–4.48 (m, 2H), 3.90–3.35 (m, 4H), 2.33 (s, 3H), 2.02–1.74 (m, 4H), 1.44 (s, 9H), 1.31 (d, *J* = 6.9 Hz, 3H); ESI MS m/z 363 [C20H30N2O4 + H]+.

#### Preparation of (*S*)-2-Amino-1-(4-(*m*-tolyloxy)piperidin-1-yl)propan-1-one

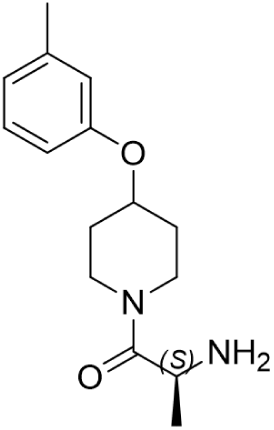

To a stirred solution of *tert*-butyl (*S*)-(1-oxo-1-(4-(*m*-tolyloxy)piperidin-1-yl)propan-2-yl)carbamate (4.53 g, 12.5 mmol) in dichloromethane (38 mL) precooled in an ice-bath was added 2,2,2-trifluoroacetic acid (9.60 mL, 125 mmol) dropwise. The resulting mixture was stirred for 21 h and left to slowly warm to room temperature. After this time, the reaction mixture was concentrated, re-dissolved in dichloromethane (125 mL) and washed with 1:1 saturated aqueous sodium bicarbonate/water (2 × 100 mL), and brine (100 mL). The aqueous layers were combined and extracted with dichloromethane (50 mL). The combined organics were dried over sodium sulfate, the solids removed by filtration, and the solvents removed *in vacuo* to afford (*S*)-2-amino-1-(4-(*m*-tolyloxy)piperidin-1-yl)propan-1-one (3.04 g, 93%) as a light yellow oil which was used without further purification: ¹H NMR (300 MHz, CDCl3) δ 7.17 (t, *J* = 7.8 Hz, 1H), 6.82–6.68 (m, 3H), 4.61–4.49 (m, 1H), 3.94– 3.59 (m, 4H), 3.53–3.32 (m, 1H), 2.33 (s, 3H), 2.26 (s, 2H), 2.03–1.73 (m, 4H), 1.28 (d, *J* = 6.9 Hz, 3H); ESI MS m/z 263 [C15H22N2O2 + H]+.

#### Preparation of (*S*)-1-(4-(*m*-Tolyloxy)piperidin-1-yl)propan-2-amine

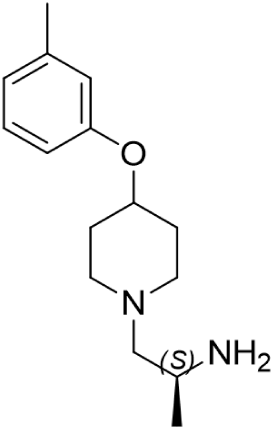

To a stirred room temperature solution of (*S*)-2-amino-1-(4-(*m*-tolyloxy)piperidin-1-yl)propan-1-one (5.23 g, 19.9 mmol) in tetrahydrofuran (250 mL) under a nitrogen atmosphere was added borane dimethyl sulfide complex (9.46 mL, 100 mmol) dropwise. The reaction mixture was then placed in a pre-heated heating block (65 °C) and stirred for 4 h. After this time, the reaction mixture was cooled in an ice-bath and stirred for 10 min before slowly adding methanol (24.2 mL, 0.598 mmol) in 1–2 mL portions. (*Caution: Vigorous gas evolution occurred during the addition of the first 6–8 mL of methanol*). The resulting mixture was stirred for 5 min, the ice-bath was then removed, and the resulting mixture was stirred at room temperature for 1 h. After this time, hydrochloric acid (1 M solution in diethyl ether; 49.8 mL, 49.8 mmol) was added and the reaction was reheated to 65 °C for 1.5 h. The reaction mixture was cooled to room temperature using a water bath, further cooled using an ice-bath, and then quenched with 1 M aqueous sodium hydroxide (∼100 mL) to pH ∼10. After stirring for 5 min, the ice-bath was removed and the reaction mixture diluted with ethyl acetate (200 mL) and water (200 mL). The organic layer was separated, and the aqueous layer was extracted with ethyl acetate (100 mL). The combined organic layer was dried over sodium sulfate, the solids removed by filtration and the solvents removed *in vacuo*. The resulting crude material was then purified by column chromatography (120 g, silica, 5 to 80% CMA (80% chloroform, 18% methanol, 2% ammonium hydroxide)/methylene chloride) to afford (*S*)-1-(4-(*m*-tolyloxy)piperidin-1-yl)propan-2-amine (3.71 g, 75%) as a clear oil: ¹H NMR (300 MHz, CDCl3) δ 7.15 (t, *J* = 7.7 Hz, 1H), 6.73 (t, *J* = 9.5 Hz, 3H), 4.36–4.21 (m, 1H), 3.12–2.96 (m, 1H), 2.91–2.76 (m, 1H), 2.72–2.57 (m, 1H), 2.47–2.34 (m, 1H), 2.32 (s, 3H), 2.26–2.06 (m, 3H), 2.05–1.90 (m, 2H), 1.88–1.70 (m, 2H), 1.70–1.60 (s, 2H), 1.02 (d, *J* = 6.3 Hz, 3H); ESI MS m/z 249 [C15H24N2O + H]+.

#### Preparation of (*S*)-4-(((1-(4-(*m*-Tolyloxy)piperidin-1-yl)propan-2-yl)amino)methyl)pyridine 1-Oxide Trihydrochloride

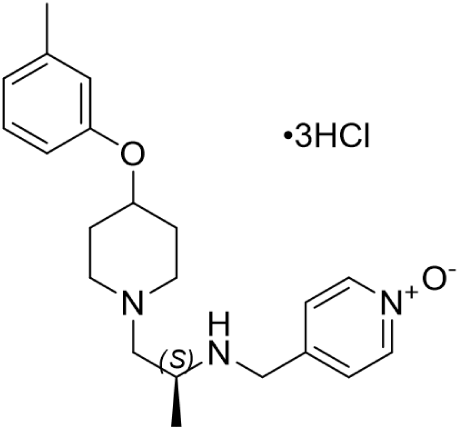

To a stirred room temperature solution of (*S*)-1-(4-(*m*-tolyloxy)piperidin-1-yl)propan-2-amine (3.70 g, 14.9 mmol) in anhydrous dichloromethane (111 mL) under a nitrogen atmosphere was added 4-formylpyridine 1-oxide (1.83 g, 14.9 mmol), magnesium sulfate (17.9 g, 149 mmol), and pre-dried 4 Å powdered molecular sieves (2.6 g). The resulting reaction mixture was stirred for 15 h. After this time, the reaction mixture was then filtered through a celite plug and washed with dichloromethane (2 × 50 mL). The combined filtrate was concentrated *in vacuo*, and the resulting reaction residue was dissolved in anhydrous methanol (111 mL). To the stirred room temperature solution was added sodium borohydride (0.620 g, 16.4 mmol) portion wise. After stirring for 1 h under a nitrogen atmosphere, the reaction mixture was diluted with saturated aqueous sodium bicarbonate solution (150 mL), water (200 mL), and ethyl acetate (150 mL). The organic layer was separated, and the aqueous layer was extracted with ethyl acetate (4 × 150 mL). The combined organic layer was dried over sodium sulfate, the solids removed by filtration, and the solvents removed *in vacuo*. The crude material was initially purified by column chromatography (120 g, silica gel, 0 to 35% methanol/methylene chloride over 35 min) to afford impure product. The fractions containing desired compound were combined and concentrated *in vacuo*. The impure desired compound was then re-purified by column chromatography (40 g, silica gel, 0 to 25% CMA (80% chloroform, 18% methanol, 2% ammonium hydroxide)/methylene chloride over 10 min, the gradient was held for 5 min, and then 25 to 45% CMA/methylene chloride over 5 min) to afford pure product as a free base. The free base (∼3 g) was dissolved in acetonitrile (30 mL) and concentrated *in vacuo.* This was repeated twice. The resulting residue was dissolved in acetonitrile (10 mL) and 2 M aqueous hydrochloric acid (30 mL) was added, resulting in a slightly cloudy solution. Water (10 mL) was added and the resulting mixture was lyophilized to afford (*S*)-4-(((1-(4-(*m*-tolyloxy)piperidin-1-yl)propan-2-yl)amino)methyl)pyridine 1-oxide trihydrochloride (3.71 g, 53%) as a white solid: ¹H NMR (300 MHz, DMSO-*d6*) δ 10.88 (br s, 1H), 10.03 (br s, 1H), 9.77 (br s, 1H), 8.37 (d, *J* = 6.6 Hz, 2H), 7.74 (d, *J* = 6.6 Hz, 2H), 7.25–7.13 (m, 1H), 6.91–6.71 (m, 3H), 4.87–2.98 (m, 11H), 2.39–1.83 (m, 7H), 1.48 (d, *J* = 6.0 Hz, 3H); ESI MS m/z 356 [C21H29N3O2 + H]+. UPLC (Method A), tR = 2.42 min, >99% (AUC) at 254 nm and >99% (AUC) at 215 nm.

Elemental analysis: calculated for C21H29N3O2•3HCl•0.1H2O: C, 54.05; H, 6.95; N, 9.00; Cl, 22.79; found: C, 54.15; H, 6.94; N, 8.90; Cl, 22.43. For *in vitro* experiments, C273 was prepared as a 10 mM stock solution in DMSO. Stock aliquots of 20 µl were stored at -20 °C.

### Mitochondrial isolation and measurement of electron transport chain (ETC) complex activity

Intact brain mitochondria were isolated from mouse brain tissue via differential centrifugation with digitonin treatment^28^. The brain tissue was immersed in ice-cold isolation medium (225 mM mannitol, 75 mM sucrose, 20 mM HEPES-Tris, and 1 mM EGTA, pH 7·4) supplemented with 1 mg/ml BSA. The tissue was homogenized with 40 strokes by the “B” (tight) pestle of a Dounce homogenizer in 10 ml of isolation medium, diluted twofold and transferred into centrifuge tubes. The homogenate was centrifuged at 5,900 × g for 4 min in a refrigerated (4 °C) Beckman centrifuge. The supernatant was centrifuged at 12,000 × g for 10 min, the pellets were resuspended in the same buffer, and 0.02% digitonin was added. The suspension was homogenized briefly with five strokes in a loosely fitted Potter homogenizer and centrifuged again at 12,000 × g for 10 min, then gently resuspended in isolation buffer without BSA and washed once by centrifuging at 12,000 × g for 10 min. The final mitochondrial pellet was resuspended in 0.1 ml of washing buffer and stored on ice. The activity of mtCI was measured spectrophotometrically via a plate reader (SpectraMax M5, Molecular Devices, USA) in 0.2 ml of standard respiration buffer composed of 125 mM sucrose, 25 mM Tris-HCl (pH = 7·5), 0.01 mM EGTA, and 20 mM KCl at 25 °C. The NADH-dependent activity of complex I was assayed as oxidation of 0.15 mM NADH at 340 nm (ε340 nm = 6·22 mM-1 cm-1) in assay buffer supplemented with 10 µM cytochrome c, 40 µg/ml alamethicin, and 1 mM MgCl_2_ (NADH media). NADH:Q reductase activity was measured in NADH media containing 2 mg/ml BSA, 60 µM decylubiquinone, 1 mM cyanide and 5–15 µg protein per well. The activity of the ETC complexes was measured via complex I (Cayman Chemicals, cat. #700930), complex II/III (Cayman Chemicals, cat. # 700950), complex IV (Cayman Chemicals, cat. # 700990), and complex V (Cayman Chemicals, cat. # 701000) colorimetric assay kits.

### Real-time respirometry

The kinetic injection experiment was performed via the Agilent Seahorse XFe96 Extracellular Flux Analyzer. Prior to the assay, the media was replaced with Agilent Seahorse XF DMEM, pH 7.4 (Agilent, 103575-100), supplemented with 1 mM pyruvate, 2 mM glutamine, and 10 mM glucose at 37 °C for 1 hour and placed in a BioTek Cytation 5 Cell Imaging Multimode Reader (Agilent). During the assay run, the compounds were injected via one of the ports, and the oxygen consumption rate (OCR) was measured every 18 minutes after compound injection for 1 hour.

### Mitochondrial stress test

This assay was performed to determine mitochondrial bioenergetics. Before each assay, the media was exchanged for Agilent Seahorse XF DMEM pH 7.4 (Agilent, 103575-100) supplemented with 1 mM pyruvate, 2 mM glutamine, and 10 mM glucose at 37 °C for 1 hour without CO_2_. Additionally, brightfield images were obtained via a BioTek Cytation 5 Cell Imaging Multimode Reader (Agilent). The oxygen consumption rate (OCR) was analysed under basal conditions and after treatment with different drugs, including the ATP synthase inhibitor oligomycin A (the optimal dose chosen after the dose‒response optimization assay: 0.5–3 µM oligomycin A), an ETC uncoupler FCCP (the optimal dose chosen after the dose‒response optimization assay: 0.5–3 µM FCCP), and an ETC inhibitor mixture (0.5 µM rotenone and 0.5 µM antimycin A) (Seahorse XF Cell Mito Stress Test Kit, Agilent, 103015-100). The response to the minimal dose of oligomycin A (2 µM) and FCCP (2 µM), generating the maximal effect, accounts for nonphosphorylating mitochondrial respiration and maximal FCCP-uncoupled respiration, respectively. The response to the rotenone and antimycin A mixture accounts for nonmitochondrial oxygen consumption. Hoechst 3342 dye (final concentration of 5 μM) was injected at the end of the assay, and the mixture was incubated at 37 °C for 15–30 min before fluorescence imaging via XF Imaging and Cell Counting Software for normalization to the cell count. The spare respiratory capacity (SRC) was calculated as the difference between the maximal and basal OCRs.

### Measurement of Substrate-Specific OXPHOS Activity in Permeabilized Cells Using the XFe96 Seahorse Analyzer

Activities of mitochondrial respiratory complexes were assessed in MC65 cells using the XF Plasma Membrane Permeabilizer (PMP) reagent (Agilent, Cat. #102504-100) according to the manufacturer’s protocol. XF96 cell culture microplates (Agilent, Cat. #101085-004) were coated with poly-D-lysine (12.5 μg/mL in sterile water) by adding 20 μL per well and incubating for 4 h at room temperature. Plates were then washed three times with 200 μL sterile water and allowed to dry for at least 2 h. MC65 cells were subsequently seeded at 50,000 cells per well. The day prior to the assay, the sensor cartridge was hydrated overnight in 200 μL Seahorse XF Calibrant at 37°C in a non-CO₂ incubator. Forty-eight hours after seeding, cells were treated for 2 h with vehicle or C273 (25 μM or 50 μM). Immediately prior to the assay, cell density and distribution were verified by brightfield imaging using a Cytation 5 Cell Imaging MultiMode Reader (Agilent).

Following treatment, cells were washed twice with 200 μL of 1× MAS buffer (Agilent) using rapid but gentle aspiration to minimize exposure time and preserve cell integrity. After the final wash, 180 μL of pre-warmed MAS buffer supplemented with mitochondrial substrates, ADP, and XF PMP was added to each well. For acute assessment of mitochondrial respiration, C273 (25 or 50 μM) or vehicle was maintained in the assay medium during the Seahorse run. Oxygen consumption rate (OCR) was measured using the Seahorse XFe96 Analyzer following sequential injections of defined substrates and inhibitors according to the Agilent permeabilized-cell assay workflow. Mitochondrial complex-specific activities were calculated as oxygen consumption rates under defined substrate conditions and expressed as a percentage of vehicle-treated controls.

### Cell viability, NADH, NADPH, Glutathione and Lactate assays

Cell viability was measured using the CellTiter-Glo® 2.0 assay according to the manufacturer’s instructions (Promega, WI). Intracellular NADH and NADPH levels were quantified using the NAD/NADH-Glo™ Assay (Promega, G9071) and the NADP/NADPH-Glo™ Assay (Promega, G9081), respectively. Glutathione levels were measured using the GSH/GSSG-Glo™ Assay (Promega, V6612). Lactate levels were determined using the Lactate-Glo™ Assay (Promega, J5021). Antioxidant capacity of C273 was determined using the Antioxidant Assay kit (Sigma-Aldrich, MAK334).

### *In vitro* ADME and safety studies

The *in vitro* ADME and safety profile of C273 including inhibition of major human cytochrome P450 (CYP) isoforms, hERG channel inhibition, aqueous solubility, plasma protein binding, and microsomal stability were conducted by the Contract Research Organization (CRO) Pharmaron Beijing Co., Ltd. (China). Full reports are available upon request.

### In vivo C273 pharmacokinetic studies

The in vivo pharmacokinetic (PK) and brain distribution studies of C273 were conducted in male C57BL/6J mice (8–12 weeks old) by the contract research organization Jubilant Biosys (India). Briefly, mice received either a single intravenous (IV) dose of C273 (3 mg/kg) or a single oral (PO) dose (25 mg/kg). For the IV cohort, plasma and brain samples were collected at eight time points: 0.08, 0.25, 0.5, 1, 2, 4, 8, and 24 h post-dose. For the PO cohort, samples were collected at eight time points: 0.25, 0.5, 1, 2, 4, 6, 8, and 24 h post-dose.

C273 concentrations in plasma and brain homogenates were quantified using a validated bioanalytical LC–MS/MS method. Pharmacokinetic parameters were calculated using standard noncompartmental analysis (NCA). The area under the concentration–time curve (AUC) was determined using conventional trapezoidal integration with extrapolation to infinity. A detailed study report is available upon request.

### Toxicology assessment of C273 in WT mice

*In vivo* toxicology studies were conducted in 2-month-old female wild-type (WT) mice treated with vehicle or C273 at 20, 50, or 80 mg/kg/day administered in drinking water for 30 days (n = 5 per group; IACUC protocol #A00001186-16-R24). Body weight and water consumption were monitored weekly. At the end of the treatment period, locomotor activity was assessed using the open field test. Following 30 days of treatment, mice were euthanized by cervical dislocation, and blood and tissues, including heart, liver, brain, spleen, lung, kidney, bladder, small intestine, large intestine, and eye, were collected. Tissues were fixed in 10% neutral buffered formalin for 48 h and subsequently transferred to PBS for long-term storage. Plasma and one hemisphere of the brain were flash-frozen in liquid nitrogen and stored at −80 °C.

### Histopathology

Histopathological analyses of hearts and livers collected from mice treated with vehicle or C273 (20, 50, or 80 mg/kg/day) and fixed as described above were performed by an independent pathologist blinded to treatment groups at the Department of Comparative Medicine, Mayo Clinic, Arizona. Fixed tissues were embedded in paraffin, and 4-µm sections were cut on a Leica RM2255 microtome, floated on a warm water bath, and mounted onto positively charged glass slides. Paraffin-embedded sections were stained with hematoxylin (Fisher, catalog #22-050-112) and eosin (Cardinal Health, Epredia series, catalog #7111) using standard H&E techniques on a Leica ST5010 autostainer and coverslipped on a Leica CV5030. Stained slides were reviewed on an Olympus BX45 microscope by a board-certified veterinary pathologist (N.M.G.), and photomicrographs were acquired with an Olympus SC180 camera (scale bars, 500 µm).

### Western blot analysis

Protein levels in the cortico-hippocampal region of the brains of vehicle- or C273-treated C57BL/6J female mice were determined via Western blot analysis. The tissue was homogenized and lysed via RIPA buffer (25 mM Tris-HCl pH 7.6, 150 mM NaCl, 1% NP-40, 1% sodium deoxycholate, 0.1% SDS) containing phosphatase PhosSTOP (Roche, cat. #04906837001) and protease inhibitors (cOmplete, Roche, cat. #11697498001). Total protein lysates (25 μg) were separated on 4–20% Mini-PROTEAN TGX™ Precast Protein Gels (Bio-Rad, 4561093). For the C273 time course study, coronal brain slices that encompassed the cortico-hippocampal region were homogenized, and 30 µg of protein lysates were separated on 4–15% Criterion gels (Bio-Rad, cat. # 5678083) and transferred to Immun-Blot polyvinylidene difluoride membranes (PVDF cat. # 1620177). Total cell lysates were prepared using RIPA buffer. Bands were imaged and quantified using ChemiDoc™ MP Imaging System (Bio-Rad, USA).

### Statistics

All the statistical analyses were performed via GraphPad Prism 10.5.0. The statistical analysis included two-tailed unpaired and paired Student’s *t* tests (where appropriate),ordinary one-way ANOVA and two-way ANOVA. The power calculation was based on a one-way or two-way ANOVA comparing up to four groups, assuming a power of 0.7, a large effect size (Cohen’s f ≈ 0.4, equivalent to d ≈ 0.75), and a significance level of 0.05. Based on these parameters, a minimum of five animals *per* group was estimated. The study was not powered to assess sex-specific differences. When *P* values were significant at a level of *P* < 0.05, Dunnett and Fisher’s LSD *post hoc* analysis for the multiple comparisons were applied to determine the differences among groups. The data are presented as the means ± SDs for each group of mice.

## Data availability

All data, including raw datasets generated or analyzed during this study, are included in this published article and its Supplementary Materials/Information file. The C273 structure was submitted in PubChem with submission number 135463. All data regarding the experiments described in the paper are available upon request to the corresponding author Eugenia Trushina.

## Acknowledgements

This research was supported by grants from NIH R01AG 55549 and UG3/UH3 NS 113776 (all to ET); U19 AG069701 (to TK); the Alzheimer’s Association Research Fellowship grant 23AARF-1027342 (to TKON). We thank members of the Research Histology and Pathology Core at Mayo Clinic in Arizona, Ms. J. Pattengill, Mrs. B. Palmer and S. Lesueur, Dr. N. M. Gades, for help with tissue pathology; Ms. S. Gochnauer for help with the manuscript preparation; former and current members of Dr. Trushina lab for their support. We thank CROs Curia, Charnwood Molecular, Pharmaron, Jubiliant Biosys for assistance with compound synthesis, medicinal chemistry efforts, ADME, PK and other tests. This research is solely the responsibility of the authors and does not necessarily represent the official view of the NIH. The funders had no role in the study design, data collection and analysis, decision to publish, or preparation of the manuscript.

## Contributions

E.T. conceived the study, assembled the multidisciplinary team of collaborators, and received funding for the project. S.T., T.K.O.N., M.O., and A.G. performed the experiments and analysed and interpreted the data. G.J. supervised medicinal chemistry efforts, T.K.O.N., T.N., W.L. and T.K. conducted the experiments on the human organoids, and S.T. and E.T. wrote the manuscript. All authors read and approved the final version of the manuscript. E.T. and S.T. directly assessed and verified the underlying data presented in the manuscript.

## Ethics declaration

ET, MO, ST and TKON are coauthors on patents relevant to the development of small molecules mitochondrial modulators. The authors declare no competing interests. The manuscript was professionally edited by Rubriq. No portion was generated using AI.

## Supplementary Figures Legends

**Figure S1. Kinetic injection ECAR data, hERG inhibition and lactate accumulation.** a The Extracellular Acidification Rate (ECAR) was measured following the injection of 25, 50 and 100 μM of C273 using the Agilent Seahorse XFe96 Extracellular Flux Analyzer. b hERG inhibition of C273 in Dofetilide radioligand binding assay performed by CRO Pharmaron. IC_50_ was generated using GraphPad Prism 10. c Lactate levels in C273-treated hepatic HepG2 cell line was measured using luminescent Lactate-Glo^TM^ assay(Promega).

**Figure S2. Western blot quantification for Fig. 4e**. Bands were imaged and quantified using ChemiDoc™ MP Imaging System (Bio-Rad, USA).

**Figure S3. Raw Western blot images for Fig. 4e**. Bands were imaged and quantified using ChemiDoc™ MP Imaging System (Bio-Rad, USA).

**Figure S4. Western blot quantification for Fig. 5a**. Bands were imaged and quantified using ChemiDoc™ MP Imaging System (Bio-Rad, USA).

**Figure S5. Raw Western blot images for Fig. 5a**. Bands were imaged and quantified using ChemiDoc™ MP Imaging System (Bio-Rad, USA).

**Figure S6. C273 does not exhibit direct antioxidant activity and inhibits TNFα-induced NF-κB activity. a** Antioxidant capacity was assessed using the Total Antioxidant Capacity Assay (Sigma). b NF-**κ**B reporter HEK293 cells were pretreated for 24 h with C273 (50 nM, 500 nM, or 5 μM), AICAR (0.5 mM), or metformin (0.5 mM), followed by stimulation with TNFα (5 ng/ml) for 6 h. NF-**κ**B Luciferase activity was measured using the Luciferase Assay System (Promega, E1501).

**Figure S7. Raw Western blot images for Fig. 6e**.

**Figure S8. Western blot quantification for Fig. 6e** Bands were imaged and quantified using ChemiDoc™ MP Imaging System.

**Figure S9. Design of C273 safety study. a Timeline of C273 treatment in WT mice b Groups, treatment and weights of mice.**

**Figure S10. Raw Western blot images for Fig. 8a. Figure S11. Raw Western blot images for Fig. 8b**.

**Figure S12. Raw Western blot images and quantification for Fig. 8c**.

**Figure S13. iPSC organoid data-Levels of Aβ40, Aβ42, p-Tau and t-Tau. a** The overview of the procedures for generating human iPSC-derived organoids from LOAD patients. b-e Levels of Aβ40 and Aβ42 in the RIPA lysate from human iPSC-derived organoids treated with vehicle, CP2, or C273 for 48 h were measured using ELISA. f-i Levels of total Tau and p-Tau (T181) in the RIPA lysate from human iPSC-derived organoids treated with vehicle, CP2, or C273 for 48 h were measured using ELISA. Data were normalized to the total protein concentration of each sample (*n* = 3 organoids *per* treatment, measured in duplicates). Data are expressed as mean ± SD. Statistical analysis was performed using a nonparametric one-way ANOVA.

**Figure S14. iPSC organoid data-Quantification of Western blots and raw images. a,b** Quantification of Western blots for pAMPKα and pGSK3β shown on Fig. 8f,g. c Quantification of p-Tau (T181) to t-Tau46 shown on Fig. 8f,g. d Raw images of Western blots shown on Fig. 8 f,g.

**Figure S15. Raw images of p-Tau (T181) to t-Tau46 are shown in Fig. 8 f,g.**

**Figure S16. iPSC organoid data-trial 2. a,b** Western blots of human iPSC-derived organoids treated with vehicle, CP2 (250 nM and 500 nM) or C273 (500 nM and 1000 nM) (second trial). c-h Raw images of Western blots shown on Figure S16 a,b.

Supplementary Table 1. CEREP44 Safety panel for C273.

## References

1 Knopman, D. S. et al. Alzheimer disease. Nat Rev Dis Primers 7, 33 (2021). 10.1038/s41572-021-00269-y

2 Zheng, Q. & Wang, X. Alzheimer’s disease: insights into pathology, molecular mechanisms, and therapy. Protein Cell 16, 83–120 (2025). 10.1093/procel/pwae026

3 Xiao, X., Yan, X., Liang, C. & Yang, Y. Metabolic dysfunction and mitochondrial failure in Alzheimer’s disease: integrating pathophysiology, clinical evidence and emerging interventions. Front Neurol 17, 1772036 (2026). 10.3389/fneur.2026.1772036

4 Ashleigh, T., Swerdlow, R. H. & Beal, M. F. The role of mitochondrial dysfunction in Alzheimer’s disease pathogenesis. Alzheimers Dement 19, 333–342 (2023). 10.1002/alz.12683

5 Wang, W., Zhao, F., Ma, X., Perry, G. & Zhu, X. Mitochondria dysfunction in the pathogenesis of Alzheimer’s disease: recent advances. Mol Neurodegener 15, 30 (2020). 10.1186/s13024-020-00376-6

6 Caberlotto, L. et al. Cross-disease analysis of Alzheimer’s disease and type-2 Diabetes highlights the role of autophagy in the pathophysiology of two highly comorbid diseases. Sci Rep 9, 3965 (2019). 10.1038/s41598-019-39828-5

7 Jurcau, M. C. et al. The Link between Oxidative Stress, Mitochondrial Dysfunction and Neuroinflammation in the Pathophysiology of Alzheimer’s Disease: Therapeutic Implications and Future Perspectives. Antioxidants (Basel) 11 (2022). 10.3390/antiox11112167

8 Li, X. et al. The role of mitochondrial dysfunction in the pathogenesis of Alzheimer’s disease and future strategies for targeted therapy. Eur J Med Res 30, 434 (2025). 10.1186/s40001-025-02699-w

9 Eckert, A., Schmitt, K. & Gotz, J. Mitochondrial dysfunction - the beginning of the end in Alzheimer’s disease? Separate and synergistic modes of tau and amyloid-beta toxicity. Alzheimers Res Ther 3, 15 (2011). 10.1186/alzrt74

10 Trushina, E., Trushin, S. & Hasan, M. F. Mitochondrial complex I as a therapeutic target for Alzheimer’s disease. Acta Pharm Sin B 12, 483–495 (2022). 10.1016/j.apsb.2021.11.003

11 Hirst, J. Mitochondrial complex I. Annu Rev Biochem 82, 551–575 (2013). 10.1146/annurev-biochem-070511-103700

12 Weinberg, S. E. & Chandel, N. S. Mitochondria reactive oxygen species signaling-dependent immune responses in macrophages and T cells. Immunity 58, 1904–1921 (2025). 10.1016/j.immuni.2025.07.012

13 Santidrian, A. F. et al. Mitochondrial complex I activity and NAD+/NADH balance regulate breast cancer progression. J Clin Invest 123, 1068–1081 (2013). 10.1172/JCI64264

14 Meichsner, A., Bader, V. & Winklhofer, K. F. Mitochondria as sources and targets of cellular signaling. Mol Cell 86, 503–521 (2026). 10.1016/j.molcel.2026.01.008

15 Chandel, N. S. Evolution of Mitochondria as Signaling Organelles. Cell Metab 22, 204–206 (2015). 10.1016/j.cmet.2015.05.013

16 Rose, G., Santoro, A. & Salvioli, S. Mitochondria and mitochondria-induced signalling molecules as longevity determinants. Mech Ageing Dev 165, 115–128 (2017). 10.1016/j.mad.2016.12.002

17 Sherer, T. B. et al. Mechanism of toxicity in rotenone models of Parkinson’s disease. J Neurosci 23, 10756–10764 (2003). 10.1523/JNEUROSCI.23-34-10756.2003

18 Bennett, C. F., Latorre-Muro, P. & Puigserver, P. Mechanisms of mitochondrial respiratory adaptation. Nat Rev Mol Cell Biol 23, 817–835 (2022). 10.1038/s41580-022-00506-6

19 Spaulding, H. R. & Yan, Z. AMPK and the Adaptation to Exercise. Annu Rev Physiol 84, 209–227 (2022). 10.1146/annurev-physiol-060721-095517

20 Steinberg, G. R. & Hardie, D. G. New insights into activation and function of the AMPK. Nat Rev Mol Cell Biol 24, 255–272 (2023). 10.1038/s41580-022-00547-x

21 Herzig, S. & Shaw, R. J. AMPK: guardian of metabolism and mitochondrial homeostasis. Nat Rev Mol Cell Biol 19, 121–135 (2018). 10.1038/nrm.2017.95

22 Jager, S., Handschin, C., St-Pierre, J. & Spiegelman, B. M. AMP-activated protein kinase (AMPK) action in skeletal muscle via direct phosphorylation of PGC-1αlpha. Proc Natl Acad Sci U S A 104, 12017–12022 (2007). 10.1073/pnas.0705070104

23 Rabinovitch, R. C. et al. AMPK Maintains Cellular Metabolic Homeostasis through Regulation of Mitochondrial Reactive Oxygen Species. Cell Rep 21, 1–9 (2017). 10.1016/j.celrep.2017.09.026

24 Salminen, A., Hyttinen, J. M. & Kaarniranta, K. AMP-activated protein kinase inhibits NF-kappaB signaling and inflammation: impact on healthspan and lifespan. J Mol Med (Berl) 89, 667–676 (2011). 10.1007/s00109-011-0748-0

25 Stojakovic, A. et al. Partial Inhibition of Mitochondrial Complex I Reduces Tau Pathology and Improves Energy Homeostasis and Synaptic Function in 3xTg-AD Mice. J Alzheimers Dis 79, 335–353 (2021). 10.3233/JAD-201015

26 Stojakovic, A. et al. Partial inhibition of mitochondrial complex I ameliorates Alzheimer’s disease pathology and cognition in APP/PS1 female mice. Commun Biol 4, 61 (2021). 10.1038/s42003-020-01584-y

27 Trushin, S. et al. Therapeutic assessment of a novel mitochondrial complex I inhibitor in in vitro and in vivo models of Alzheimer’s disease. EBioMedicine 120, 105924 (2025). 10.1016/j.ebiom.2025.105924

28 Zhang, L. et al. Modulation of mitochondrial complex I activity averts cognitive decline in multiple animal models of familial Alzheimer’s Disease. EBioMedicine 2, 294–305 (2015). 10.1016/j.ebiom.2015.03.009

29 Gao, H. et al. A genome-wide association study in human lymphoblastoid cells supports safety of mitochondrial complex I inhibitor. Mitochondrion 58, 83–94 (2021). 10.1016/j.mito.2021.02.005

30 Gao, H. et al. Mitochondrial complex I deficiency induces Alzheimer’s disease-like signatures that are reversible by targeted therapy. Alzheimers Dement 21, e70519 (2025). 10.1002/alz.70519

31 Panes, J. et al. Partial Inhibition of Complex I Restores Mitochondrial Morphology and Mitochondria-ER Communication in Hippocampus of APP/PS1 Mice. Cells 12 (2023). 10.3390/cells12081111

32 Keller, N., Christensen, T. A., Wanberg, E. J., Salisbury, J. L. & Trushina, E. Neuroprotective mitochondria targeted small molecule restores synapses and the distribution of synaptic mitochondria in the hippocampus of APP/PS1 mice. Sci Rep 15, 6528 (2025). 10.1038/s41598-025-90925-0

33 Maezawa, I. et al. The Anti-Amyloid-beta and Neuroprotective Properties of a Novel Tricyclic Pyrone Molecule. J Alzheimers Dis 58, 559–574 (2017). 10.3233/JAD-161175

34 Trushina, E., Nguyen, T. K. O. & Trushin, S. Modulation of Mitochondrial Function as a Therapeutic Strategy for Neurodegenerative Diseases. J Prev Alzheimers Dis 10, 675–685 (2023). 10.14283/jpad.2023.108

35 Maezawa, I. et al. A novel tricyclic pyrone compound ameliorates cell death associated with intracellular amyloid-beta oligomeric complexes. J Neurochem 98, 57–67 (2006). 10.1111/j.1471-4159.2006.03862.x

36 Rankovic, Z. CNS drug design: balancing physicochemical properties for optimal brain exposure. J Med Chem 58, 2584–2608 (2015). 10.1021/jm501535r

37 Sopher, B. L. et al. Cytotoxicity mediated by conditional expression of a carboxyl-terminal derivative of the beta-amyloid precursor protein. Brain Res Mol Brain Res 26, 207–217 (1994). 10.1016/0169-328x(94)90092-2

38 Huang, L. et al. Intracellular amyloid toxicity induces oxytosis/ferroptosis regulated cell death. Cell Death Dis 11, 828 (2020). 10.1038/s41419-020-03020-9

39 Marchetti, P., Fovez, Q., Germain, N., Khamari, R. & Kluza, J. Mitochondrial spare respiratory capacity: Mechanisms, regulation, and significance in non-transformed and cancer cells. FASEB J 34, 13106–13124 (2020). 10.1096/fj.202000767R

40 Zhao, J. et al. APOE4 exacerbates synapse loss and neurodegeneration in Alzheimer’s disease patient iPSC-derived cerebral organoids. Nat Commun 11, 5540 (2020). 10.1038/s41467-020-19264-0

41 Fandos, N. et al. Plasma Abeta42/Abeta40 determined by mass spectrometry is associated with longitudinal changes in amyloid accumulation, brain atrophy, and conversion to mild cognitive impairment due to Alzheimer’s disease in individuals with subjective cognitive decline: 5-year follow-up of the FACEHBI cohort. J Prev Alzheimers Dis 13, 100465 (2026). 10.1016/j.tjpad.2025.100465

42 Chakraborty, P. et al. GSK3βeta phosphorylation catalyzes the aggregation of tau into Alzheimer’s disease-like filaments. Proc Natl Acad Sci U S A 121, e2414176121 (2024). 10.1073/pnas.2414176121

43 Sayas, C. L. & Avila, J. GSK-3 and Tau: A Key Duet in Alzheimer’s Disease. Cells 10 (2021). 10.3390/cells10040721

44 Jeremic, D., Jimenez-Diaz, L. & Navarro-Lopez, J. D. Past, present and future of therapeutic strategies against amyloid-beta peptides in Alzheimer’s disease: a systematic review. Ageing Res Rev 72, 101496 (2021). 10.1016/j.arr.2021.101496

45 Cummings, J., Osse, A. M. L., Cammann, D., Powell, J. & Chen, J. Anti-Amyloid Monoclonal Antibodies for the Treatment of Alzheimer’s Disease. BioDrugs 38, 5–22 (2024). 10.1007/s40259-023-00633-2

46 Cummings, J. L. et al. Alzheimer’s disease drug development pipeline: 2025. Alzheimers Dement (N Y) 11, e70098 (2025). 10.1002/trc2.70098

47 Papaliagkas, V. Anti-Amyloid Therapies for Alzheimer’s Disease: Progress, Pitfalls, and the Path Ahead. Int J Mol Sci 26 (2025). 10.3390/ijms26199529

48 Song, T. et al. Mitochondrial dysfunction, oxidative stress, neuroinflammation, and metabolic alterations in the progression of Alzheimer’s disease: A meta-analysis of in vivo magnetic resonance spectroscopy studies. Ageing Res Rev 72, 101503 (2021). 10.1016/j.arr.2021.101503

49 Wadan, A. S. et al. Mitochondrial-based therapies for neurodegenerative diseases: a review of the current literature. Naunyn Schmiedebergs Arch Pharmacol 398, 11357–11386 (2025). 10.1007/s00210-025-04014-0

50 Molina, J. R. et al. An inhibitor of oxidative phosphorylation exploits cancer vulnerability. Nat Med 24, 1036–1046 (2018). 10.1038/s41591-018-0052-4

51 Pujalte-Martin, M. et al. Targeting cancer and immune cell metabolism with the complex I inhibitors metformin and IACS-010759. Mol Oncol 18, 1719–1738 (2024). 10.1002/1878-0261.13583

52 Yang, Y. et al. Metformin decelerates aging clock in male monkeys. Cell 187, 6358–6378 e6329 (2024). 10.1016/j.cell.2024.08.021

53 Kulkarni, A. S., Gubbi, S. & Barzilai, N. Benefits of Metformin in Attenuating the Hallmarks of Aging. Cell Metab 32, 15–30 (2020). 10.1016/j.cmet.2020.04.001

54 Trushina, E., Rana, S., McMurray, C. T. & Hua, D. H. Tricyclic pyrone compounds prevent aggregation and reverse cellular phenotypes caused by expression of mutant huntingtin protein in striatal neurons. BMC Neurosci 10, 73 (2009). 10.1186/1471-2202-10-73

55 Salminen, A., Kaarniranta, K., Haapasalo, A., Soininen, H. & Hiltunen, M. AMP-activated protein kinase: a potential player in Alzheimer’s disease. J Neurochem 118, 460–474 (2011). 10.1111/j.1471-4159.2011.07331.x

56 Cai, Z., Yan, L. J., Li, K., Quazi, S. H. & Zhao, B. Roles of AMP-activated protein kinase in Alzheimer’s disease. Neuromolecular Med 14, 1–14 (2012). 10.1007/s12017-012-8173-2

57 Wang, X., Zimmermann, H. R. & Ma, T. Therapeutic Potential of AMP-Activated Protein Kinase in Alzheimer’s Disease. J Alzheimers Dis 68, 33–38 (2019). 10.3233/JAD-181043

58 Kim, J., Yang, G., Kim, Y., Kim, J. & Ha, J. AMPK activators: mechanisms of action and physiological activities. Exp Mol Med 48, e224 (2016). 10.1038/emm.2016.16

59 Xiang, H. C. et al. AMPK activation attenuates inflammatory pain through inhibiting NF-kappaB activation and IL-1beta expression. J Neuroinflammation 16, 34 (2019). 10.1186/s12974-019-1411-x

60 Tian, Y. et al. Mitochondrial Stress Induces Chromatin Reorganization to Promote Longevity and UPR(mt). Cell 165, 1197–1208 (2016). 10.1016/j.cell.2016.04.011

61 Olivier, S., Foretz, M. & Viollet, B. Promise and challenges for direct small molecule AMPK activators. Biochem Pharmacol 153, 147–158 (2018). 10.1016/j.bcp.2018.01.049

62 Assefa, B. T., Tafere, G. G., Wondafrash, D. Z. & Gidey, M. T. The Bewildering Effect of AMPK Activators in Alzheimer’s Disease: Review of the Current Evidence. Biomed Res Int 2020, 9895121 (2020). 10.1155/2020/9895121

63 Rana, S., Hong, H. S., Barrigan, L., Jin, L. W. & Hua, D. H. Syntheses of tricyclic pyrones and pyridinones and protection of Abeta-peptide induced MC65 neuronal cell death. Bioorg Med Chem Lett 19, 670–674 (2009). 10.1016/j.bmcl.2008.12.060

64 Dietz, A. B. et al. A novel source of viable peripheral blood mononuclear cells from leukoreduction system chambers. Transfusion 46, 2083–2089 (2006). 10.1111/j.1537-2995.2006.01033.x

